# Phosphonate production by marine microbes: exploring new sources and potential function

**DOI:** 10.1101/2020.11.04.368217

**Authors:** Marianne Acker, Shane L. Hogle, Paul M. Berube, Thomas Hackl, Ramunas Stepanauskas, Sallie W. Chisholm, Daniel J. Repeta

## Abstract

Phosphonates, organic compounds with a C-P bond, constitute 20-25% of phosphorus in high molecular weight dissolved organic matter and are a significant phosphorus source for marine microbes. However, little is known about phosphonate sources, biological function, or biogeochemical cycling. Here, we determine the biogeographic distribution and prevalence of phosphonate biosynthesis potential using thousands of genomes and metagenomes from the upper 250 meters of the global ocean. Potential phosphonate producers are taxonomically diverse, occur in widely distributed and abundant marine lineages (including SAR11 and *Prochlorococcus*) and their abundance increases with depth. Within those lineages, phosphonate biosynthesis and catabolism pathways are mutually exclusive, indicating functional niche partitioning of organic phosphorus cycling in the marine microbiome. Surprisingly, one strain of *Prochlorococcus* (SB) can allocate more than 40% of its cellular P-quota towards phosphonate production. Chemical analyses and genomic evidence suggest that phosphonates in this strain are incorporated into surface layer glycoproteins that may act to reduce mortality from grazing or viral infection. Although phosphonate production is a low-frequency trait in *Prochlorococcus* populations (~ 5% of genomes), experimentally derived production rates suggest that *Prochlorococcus* could produce a significant fraction of the total phosphonate in the oligotrophic surface ocean. These results underscore the global biogeochemical impact of even relatively rare functional traits in abundant groups like *Prochlorococcus* and SAR11.

## Introduction

In nutrient-impoverished mid-ocean gyres, microbial demand for phosphorus (P) is often so high that concentrations of inorganic phosphate are drawn down to sub nanomolar levels. Under these conditions, up to half of microbial P demand is met through the uptake and metabolism of P-containing dissolved organic matter^1^. Dissolved organic phosphorus (DOP) is a complex, poorly characterized mixture of high and low molecular weight (HMW and LMW) phosphate and phosphonate esters. Phosphate esters are common in nucleic acids and lipids, and are synthesized by all marine microbes. The presence of phosphate esters in DOP is easily explained. In contrast, phosphonates, reduced P compounds with a stable, covalent C-P bond^2^, are a poorly-understood component of marine DOP, but nevertheless constitute 20-25% of the P in HMWDOP^3^.

All phosphonate production pathways are initially catalyzed through the same steps involving the enzymes PepM, Ppd, and Pdh^2^ (Figure 1A). Despite these shared catalytic steps, the chemical diversity of phosphonates is extensive and includes small bioactive metabolites as well as macromolecules such as lipids, polysaccharides, and proteins. The functional roles and macromolecular forms that phosphonates take in marine microbes remain unknown. In contrast, phosphonate degradation potential in the marine environment is better understood^4–6^ (Figure 1B), and its distribution has been shown to be strongly shaped by phosphate availability^7,8^.

**Figure 1:**
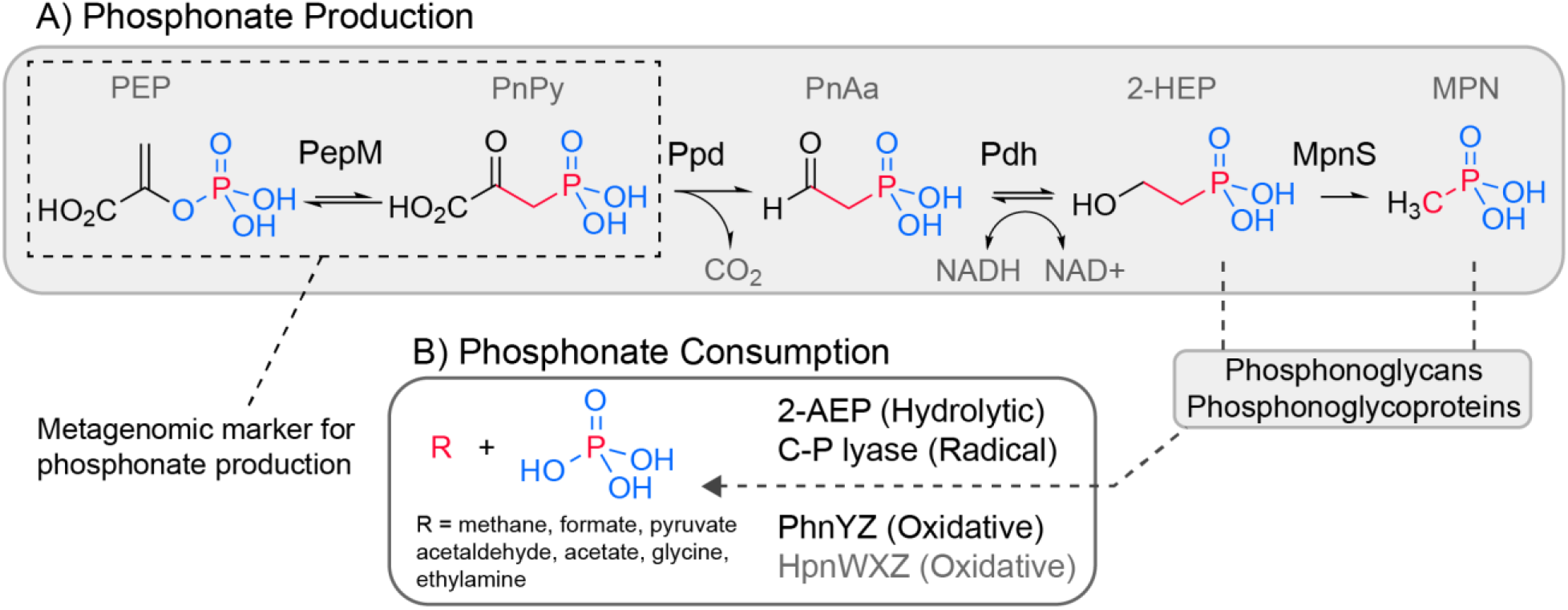
Phosphonates production and consumption in the marine environment. **A)** Phosphonate biosynthesis includes three key steps; (1) isomerization of phosphoenolpyruvate (PEP) to phosphonopyruvate (PnPy) via phosphoenolpyruvate mutase (PepM), (2) decarboxylation of PnPy to phosphonoacetaldehyde (PnAa) via phosphonopyruvate decarboxylase (Ppd), and (3) dehydrogenation of PnAa to 2-hydroxyethylphosphonic acid (2-HEP) via phosphonoacetaldehyde dehydrogenase (Pdh). 2-HEP can be further transformed by methylphosphonate synthase (MpnS), to produce methylphosphonate (MPn), a substrate for aerobic marine methane production^5^. Phosphonates in the marine environment are likely used to decorate macromolecules like glycans and glycoproteins^2,10,45^. **B)** Phosphonate degradation via cleavage of the C-P bond proceeds through at least three mechanisms: hydrolytic (2AEP, named for the representative 2-aminoethylphosphate degradation pathway via PhnWX), radical (C-P lyase), and oxidative (PhnYZ). The proposed HpnWXZ pathway is likely oxidative proceeding via the oxidative deamination of an aminophosphonate similar in structure to 2-aminoethylphosphate^106^. We find the HpnWXZ pathway to be rare in marine genomes.

Despite the high abundance of phosphonates in DOP, only two bacterioplankton (bacteria and archaea) species have been experimentally confirmed as phosphonate producers: *Trichodesmium erythraeum,* a nitrogen-fixing cyanobacterium^9^, and *Nitrosopumilus maritimus* from the Marine Group I (MGI) Thaumarchaeota^10^. *Trichodesmium* has a relatively restricted geographic range^11^ and global abundance, while MGI Thaumarcheota are abundant in the mesopelagic, but comparatively rare in sunlit surface waters^12^. Moreover, *Candidatus* Nitrosopelagicus brevis, the only characterized pelagic representative of MGI Thaumarcheota, lacks phosphonate biosynthesis genes and is significantly more abundant than *Nitrosopulimus* in the open ocean^13^. It is unlikely that phosphonate production by *Trichodesmium* and Thaumarcheota alone is enough to support the large and ubiquitous inventory of phosphonates observed in the sunlit upper ocean. Metagenomic surveys of PepM have estimated that between 8-16% of all marine microbes in the surface ocean may be capable of producing phosphonates^10,14^, but the taxonomic composition and ecological niches of phosphonate producers remain unclear.

Much about the biology, ecology, and biogeochemistry of marine phosphonates remains to be discovered. Recent vast expansions of marine genomic data^15–18^, as well as advances in chemical analyses, makes it possible to systematically investigate the question of what organisms are producing the enigmatic phosphonate pool in the oceans. Here we combine laboratory studies and chemical analyses with comparative genomic and metagenomic analyses to investigate the prevalence, taxonomic distribution, and potential function of phosphonate biosynthesis in the surface ocean. We find that phosphonate biosynthesis genes are found in a wide variety of marine microbes including the two most abundant groups in the surface ocean: *Prochlorococcus* and SAR11. We experimentally demonstrate that a *Prochlorococcus* strain produces phosphonates almost exclusively in the HMW protein fraction and, surprisingly, these phosphonoproteins account for over 40% of total cellular phosphorus.

## Main Text

### The taxonomic distribution of phosphonate producers and consumers

How widespread is the ability to produce and consume phosphonates among diverse marine microbes? Prior efforts to address this question have relied on mapping metagenomic reads to key marker genes such as the PepM found in phosphonate producers^14^ or the C-P lyase multi-enzymatic cluster and other catabolic pathway proteins found in phosphonate consumers^4,6,7^. Here we expand on prior read recruitment methods and use a genome-resolved approach with thousands of randomly sampled single-cell amplified genomes (SAGs) from the Global Ocean Reference Genome (GORG-Tropics) dataset^15^. We found that the taxonomic richness of phosphonate producers is higher than for consumers (Supplementary Note S1, Figure 2A). However, the taxonomic evenness of producers was low, with over 60% of them assigned to SAR11 clades. In contrast, phosphonate consumers were more evenly distributed, but over 90% of SAGs came from just four taxonomic orders (Figure 2B). We found that the two most numerically abundant and cosmopolitan marine groups, SAR11 and *Prochlorococcus* are likely important phosphonate producers. SAR11 are small aquatic chemoheterotrophic Alphaproteobacteria estimated to constitute up to half of the total plankton cells in the surface ocean^19^, and *Prochlorococcus* are unicellular photosynthetic picocyanobacteria that numerically dominate the euphotic zone of subtropical and tropical oligotrophic areas^20^. Although the potential for phosphonate production is broadly taxonomically distributed, highly abundant pelagic groups like SAR11 and *Prochlorococcus* are likely to be the dominant producers across subtropical/tropical surface ocean.

**Figure 2:**
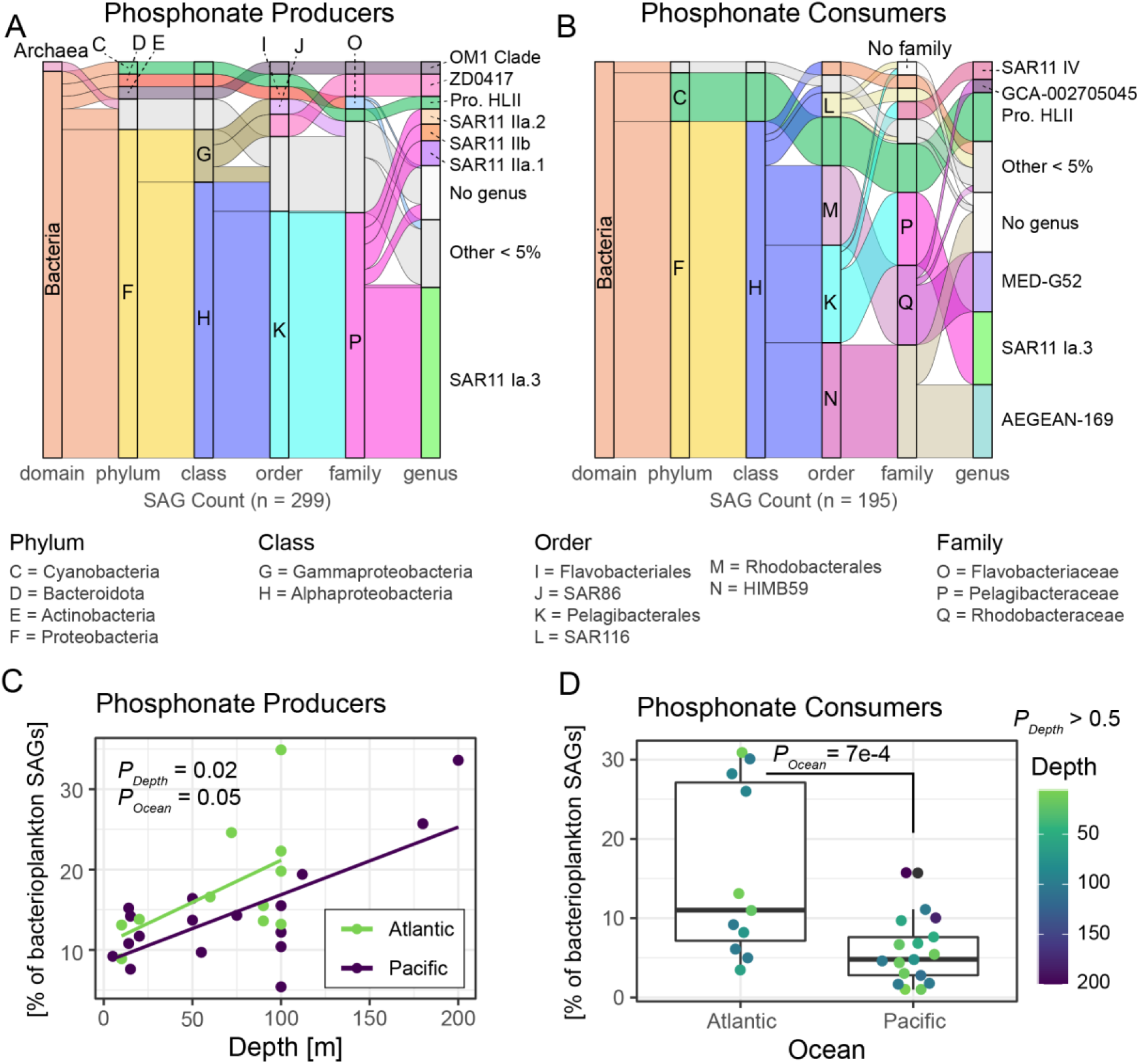
Genomic potential for phosphonate production and consumption in nearly 13000 single-cell genomes from the tropical ocean. Flow diagrams representing taxonomic hierarchy of **A)** phosphonate producers and **B)** consumers from the GORG-Tropics database15. Flows of the same color spanning multiple taxonomic ranks indicate that the higher ranks are entirely subsumed by the lowest rank. For example, all cyanobacterial genomes were from the *Prochlorococcus* genus. Otherwise each partition at each taxonomic level is colored uniquely. For the quantitative comparison of GORG-Tropics sample BATS248 was randomly subsampled to the median depth of the other 27 samples. Letters at each taxonomic rank refer to key marine archaea/bacterial groups (key left) and bar size is proportional to relative abundance. **C)** The proportion of phosphonate producers significantly increases with depth in the 28 GORG-Tropics samples. Each point is a sample (median SAGs = 241). Producers are slightly more common in the Atlantic than the Pacific [Beta-binomial regression; Depth - Est=0.0036, Err=0.0014, t=2.604, P=0.02; Ocean - Est=−0.30, Err=0.14, t=−2.08, P=0.05; link=logit; log L=−78.179, df=4, resid df=24]. **D)** Phosphonate consumers are significantly more abundant in the Atlantic than the Pacific [Beta-binomial regression; Ocean - Est=−1.04, Err=0.27, t=−3.81, P=7e-4; link=logit; log L = −80.425, df=4, resid df=24]. Proportions [%] are total producers or consumers divided by the total number GORG assemblies and are corrected using the estimated sequence recovery from assemblies (see methods).

### The abundance of phosphonate producing and consuming microbes in the oceans

We leveraged the GORG-Tropics reference database to quantitatively assess the proportion of cells in the surface ocean that are phosphonate consumers or producers. GORG-Tropics is buttressed by the use of a randomized cell selection strategy for generating SAGs – thus, minimizing issues related to functional or taxonomic biases more common to targeted genome sequencing. We estimate that, globally, 15% of all bacterioplankton in the upper 100 meters are potential phosphonate producers in the GORG-tropics database. Compared to bacteria, phosphonate producing archaea were rare in the euphotic zone (< 3% of all GORG producers) and phosphonate consuming archaea were absent (Figure 2A,B). Planktonic archaea comprised only 14% of GORG-Tropics with 80% of archaea from Marine Group II and 20% from Marine Group I. However, 37% of Marine Group I SAGs contained phosphonate biosynthesis genes. All phosphonate producing MGI archaea were isolated from below 100 meters and included both *Nitrosopumilus* (20%) and *Nitrosopelagicus* (80%). Thus, archaea may be important phosphonate producers at the base of the euphotic zone and mesopelagic, but within the sunlit ocean other bacterial producers dominate. Indeed, SAR11 (order Pelagibacterales) is by far the most abundant producer constituting over 60% of all SAGs with phosphonate biosynthesis potential (Figure 2A) and 18% of all SAR11 SAGs. Most phosphonate producers are from the surface SAR11 clades 1a.3, IIa.1 and IIa.2, and the mesopelagic IIb clade (Figure 2A, Supplementary Figure S1). The potential for methylphosphonate production is rare (< 1% of genomes with MpnS) and is predominantly found in SAR11 SAGs (80% of MpnS containing SAGs are SAR11). Metagenome recruitment from earlier studies suggested *Prochlorococcus* may account for 20% of phosphonate producers^14^. In GORG-Tropics we found that *Prochlorococcus* constitutes 6% of all phosphonate producers (Figure 2A) and 3% of all SAGs. Most *Prochlorococcus* phosphonate producers are from the surface high-light (HL) clades (Supplementary Figure S1). The percentage of bacterioplankton that can produce (15%) and consume (10%) phosphonates is similar across the global surface ocean. On average there are three times more SAR11 phosphonate producers (18%) than consumers (5%), while for *Prochlorococcus* the trend is reversed – 6% producers versus 16% consumers (Supplementary Figure S2). SAR11 has been implicated as a major contributor to marine surface water methane supersaturation due to the metabolism of methylphosphonate^21^. Here our results imply that SAR11 has an even more important global role as a phosphonate producer.

We identified two major trends for producers and consumers in GORG-Tropics: 1) the abundance of phosphonate producers significantly increases with depth and is slightly biased towards the Atlantic ocean (Figure 2C) and 2) consistent with past reports^7^ phosphonate consumers are significantly more abundant in the Atlantic ocean (Figure 2D). The same trends broadly held for SAR11 and *Prochlorococcus* consumers (Supplementary Figure S2). This motivated us to look for other relationships between phosphonate producers and environmental variables using a metagenomic dataset encompassing nearly 700 metagenomes from bioGEOTRACES, Tara Oceans, and two long-term ocean time-series sites^17,18,22^. We determined the relative abundance of the PepM in the upper 250 meters of the global ocean using read recruitment to PepM and conserved marker genes (see methods). We estimate that the median proportion of phosphonate producers in the global ocean is 6%, 10%, and 15% of *Prochlorococcus*, SAR11, and bacterioplankton genomes, respectively (Figure 3A), which agrees well with observations from GORG-Tropics. The discrepancy for SAR11 is likely explained by the low estimated sensitivity (60%) for our method to SAR11 (Supplementary Note S5). Assuming we missed 40% of SAR11 PepM sequences, the corrected fraction of SAR11 is 17% which, again, agrees well with results from GORG. There was no statistically significant difference in producer abundance between ocean basins (Supplementary Figure S3A), and no significant time-averaged difference between the Hawaii Ocean Time-series (HOT) and the Bermuda Atlantic Time Series (BATS) (Supplementary Figure S4), two long-running time series representative of the N. Pacific and N. Atlantic subtropical gyres^23,24^. However, phosphonate producers have a significant seasonal dependence at the surface at BATS where producer abundance peaks during winter (November through the following March), which coincides with the peak of wind-driven deep mixing (Supplementary Figure S4, Supplementary Note S2). Overall, the global median abundance for phosphonate producers is modest (15%) and largely stable across the global subtropical surface ocean.

**Figure 3:**
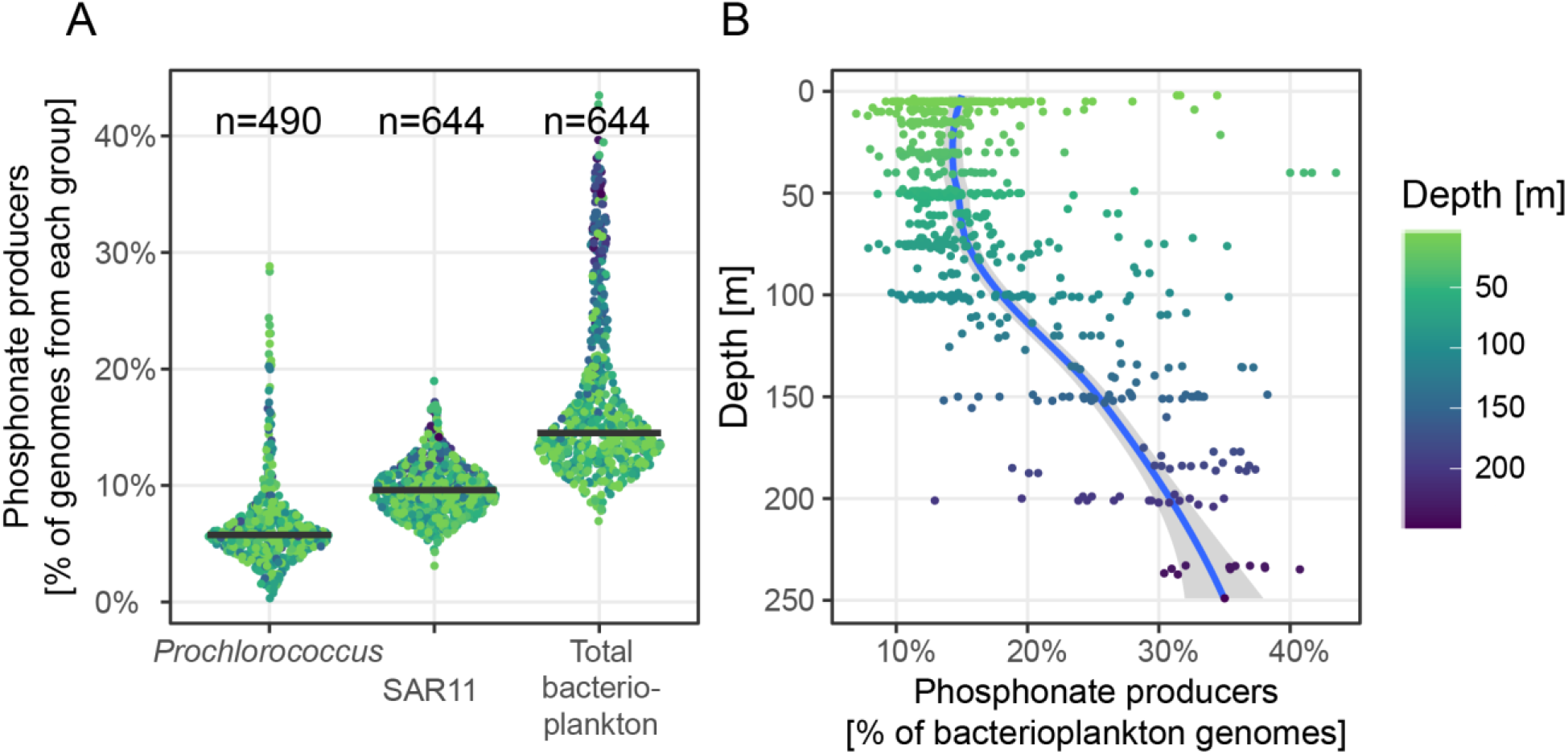
Phosphonate biosynthesis genes in surface (< 300 meters) ocean metagenomes. **A)** Phosphonate producers as the percentage of all bacterioplankton genomes estimated from combined BioGEOTRACES and Tara Oceans samples for *Prochlorococcus*, SAR11, and all Bacteria and Archaea combined. Black line is the median value for *n* metagenome samples for each taxonomic group. **B)** The relationship between depth and percent phosphonate producers in the global ocean. The blue line is a simple Loess regression fit to the data. Norm. PepM is the normalized fraction genomes with PepM and is estimated as the length-normalized abundance of PepM divided by length-normalized abundance of taxon-specific marker genes.

Are there particular ocean features that may select for the ability of a microbe to produce phosphonate? We used machine learning approaches^25^ (Supplementary Note S6) and parametric regression^26^ (Supplementary Table S1) to identify biotic and chemical/physical factors driving the distributions of *Prochlorococcus*, SAR11, and total bacterioplankton phosphonate producers (Supplementary Figure S3). Generally, we find that the greatest amount of phosphonate producer variation is explained by multiple depth-dependent biotic factors (Figure 3B, Supplementary Figure S3 and S5, Supplementary Note S3). In surface samples shallower than 100 m, a median of 15% of bacterioplankton can produce phosphonates, while below 100 m that proportion steadily increases to nearly 30-40% at 200 m. This 100 m break coincides with the nutrient-driven “genomic transition zone” i.e. a zone where bacteria and archaea tend to have larger genomes with higher GC content, and proteins with higher N/C ratios^27^, and implies that nutrient availability is an important factor setting the relative abundance of phosphonate producers. Overall it appears that a modest proportion of highly abundant oligotrophic “surface” bacterial clades including SAR11 Ia.3 and HL *Prochlorococcus* are capable of producing phosphonates in the upper 50 to 100 m and contribute to an endemic phosphonate pool there. Below 100 m, the genetic potential for phosphonate production shifts to other groups including SAR86, SAR11 IIb, the OM1 clade, MGI Archaea and the ZD0417 clade^28^. It may be that deep producers are sometimes uplifted into the upper 100 m of the euphotic zone during seasonal mixing events or ephemeral upwelling events like cyclonic mesoscale eddies^29,30^.

Our genomic and metagenomic results offer a number of predictions relevant to the biogeochemistry of marine phosphonates. First, the vertical distribution of phosphonate producers in the upper 250 meters is not even, with a distinct and rapid increase in producers beginning at 100 m depth. This implies that bulk phosphonate production rates might be similarly stratified, and that deeper phosphonate pools may be important for the total phosphonate balance in the sunlit surface ocean. Second, unlike for phosphonate consumers^7^, phosphate concentration does not appear to have a strong selective effect on the biogeography of phosphonate producers in surface waters. However, this might be due to our statistical approach not having a sufficient power to detect a true difference between ocean regions since the mean proportions we compare are small with a relatively high variability. Further investigation into ocean basin differences in phosphonate production potential is warranted. Finally, the median proportion of bacteria and archaea that can produce phosphonates in the upper 100 meters of the surface ocean appears to be relatively small (15% of bacterioplankton cells). Since phosphonates appear to be relatively labile and rapidly cycled^5^, this implies that cellular production rates would need to be quite high to account for the large global inventory of phosphonates in marine dissolved organic matter.

### Phosphonate production in Prochlorococcus SB

To better understand how much cellular P is allocated to phosphonate production and with which macromolecules they are associated, we sought to experimentally characterize phosphonate production by an abundant, widespread, and experimentally tractable marine microbe. We found only one cultured isolate that meets these criteria: *Prochlorococcus* SB. Since there is as yet no genetic system for knockout mutants in *Prochlorococcus*, we used *Prochlorococcus* MIT9301, a closely related strain lacking the phosphonate biosynthesis cluster, as a control. We analyzed cell pellets from each strain using ^31^P nuclear magnetic resonance spectroscopy (^31^P-NMR). As expected, the ^31^P-NMR spectrum of MIT9301 cells shows that all P is allocated to phosphate and pyrophosphate esters (−12 to 12 ppm; Figure 4B). In contrast, ^31^P-NMR spectrum of *Prochlorococcus* SB cells displays strong signals between both −10 and 12 ppm from phosphate and pyrophosphate esters, and between 18 and 27 ppm from phosphonates^31^ (Figure 4B). Integration of the phosphate and phosphonate ester regions of the NMR spectrum yields a cellular ratio of phosphonate to phosphate (Phn/Ph) of 0.72. (Figure 4B; Supplementary Table S3) which indicates that under nutrient replete conditions *Prochlorococcus* SB cells allocate ~ 40% of their cellular P to phosphonate production.

**Figure 4:**
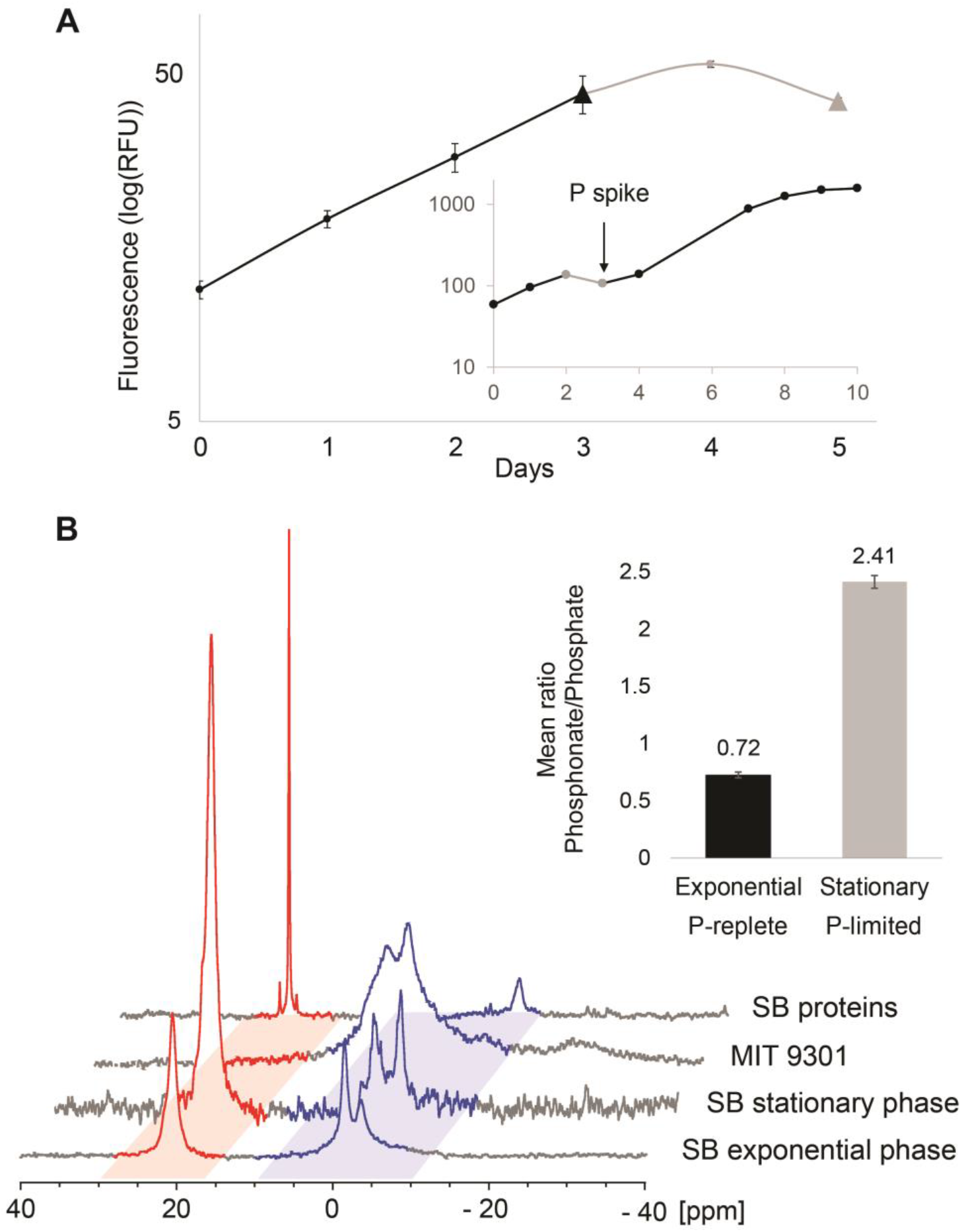
Phosphonate production by *Prochlorococcus* cultures. **A)** *Prochlorococcus* SB growth curve showing exponential phase (black) and stationary phase (grey) due to P-limitation. Triangle data points correspond to the days cultures were harvested in the different growth phases. Error bars are calculated based on the standard deviation between the biological duplicates. The inset represents the *Prochlorococcus* SB growth curve in medium with N/P = 350/1 in which inorganic phosphate was added on Day 3 to reach N/P = 16/1. **B)** ^31^P-NMR spectra of *Prochlorococcus* SB whole cells harvested in exponential phase (P-replete) and in stationary phase (P-limited), the negative control *Prochlorococcus* MIT9301 harvested in exponential phase (P-replete) and the insoluble protein fraction of *Prochlorococcus* SB harvested in exponential growth phase. The phosphonate and phosphate regions of the spectra are indicated in red and blue respectively. While *Prochlorococcus* SB produces phosphonate and doubles its relative phosphonate content in P-limited stationary phase, the negative control, MIT9301 only produces phosphates. The histogram (inset) displays the mean Phosphonate/Phosphate ratios for *Prochlorococcus* SB cells harvested in exponential (black) and stationary (grey) phase calculated by integrating the phosphonate and phosphate peaks in *Prochlorococcus* SB whole cell ^31^P-NMR spectra obtained for the duplicates in each growth phase. Error bars correspond to the standard deviation of the biological replicate phosphonate/phosphate ratio values with n=2.

### Phosphonates and P storage

The physiological and ecological roles of phosphonates are poorly understood. However, the presence of PepM among diverse genome-streamlined bacteria and the high relative abundance of cellular phosphonates in *Prochlorococcus* SB both suggest that phosphonates serve an important function for microbes inhabiting oligotrophic marine waters. One function suggested for phosphonates is that they serve as an intra-cellular P-storage reservoir^32^. In highly stratified oligotrophic waters, nutrient supply to phytoplankton is episodic; driven by mixing events that bring nutrient-rich waters from below the surface into the euphotic zone. To synchronize their need for nutrients with an episodic supply, some microbes take up excess P during periods of high nutrient concentrations and sequester them internally^33,34^. When nutrient concentrations fall, internal P-stores are metabolized to release inorganic phosphate.

The *Prochlorococcus* SB genome lacks phosphonate degradation pathways including C-P lyase, phosphonate hydrolytic pathways^2^, and PhnYZ an oxidative pathway recently discovered in *Prochlorococcus*^4^. Recognizing that *Prochlorococcus* SB might use an uncharacterized pathway to repurpose P from phosphonates, we tested the P-storage hypothesis in *Prochlorococcus* SB by comparing phosphonate production in cultures grown under P-starved and P-replete conditions (Figure 4A), expecting that if phosphonates were used for P-storage, P-limitation would reduce the allocation of P to phosphonates. However, we found that *Prochlorococcus* SB allocates significantly more P to phosphonates relative to phosphates upon entering P-limited stationary phase growth (Phn/Ph = 2.4) than during P-replete exponential growth (Phn/Ph = 0.72; Figure 4B). Cellular phosphorus to carbon (C/P) was relatively stable across exponential and stationary phases (145 and 131 respectively) and changes in Phn/Ph were driven by a decrease in phosphate ester content during P-starvation, with continued production of phosphonates (Supplementary Figure S6, Supplementary Table S4). As *Prochlorococcus* SB becomes increasingly P starved, the regulation of the cellular phosphate and phosphonate pools becomes decoupled from exponential growth conditions. While reallocation of P away from labile phosphates is one mechanism by which *Prochlorococcus* adapts to P-starvation^35^, *Prochlorococcus* SB appears to be less able to regulate phosphonate production or repurpose P in phosphonates towards other cellular functions. Both scenarios are inconsistent with phosphonates as a P-storage reservoir in *Prochlorococcus* SB, and P locked into phosphonates is not internally recycled to sustain growth during periods of P-limitation. This conclusion is reinforced by our comparative genomic analysis. Indeed, if phosphonates were used by marine microbes for luxury P storage, we would expect both phosphonate production and consumption pathways to coexist within the same genome. However, we found that genomes with phosphonate biosynthetic potential almost never encode the machinery to degrade and use phosphonates (Figure 5). In addition to biosynthesis genes, we searched for all known phosphonate degradation pathways (Figure 1). In our analysis of the GORG-Tropics and MARMICRODB^36^ genome datasets, we found that less than 1% of all genomes encoding at least one marker gene implicated in phosphonate metabolism are both producers and consumers. This mutual exclusivity occurs at the species/strain level - for example within HL *Prochlorococcus* - which implies strong functional differentiation between even closely related marine microbes. This suggests fine-scale niche partitioning between phosphonate producers and consumers in the environment and may reflect functional incompatibility or ecological/evolutionary tradeoffs between biosynthesis and catabolism. It also implies that if phosphonates were a widespread storage strategy, most phosphonate producers would have no way to reclaim P from phosphonates during times of need.

**Figure 5:**
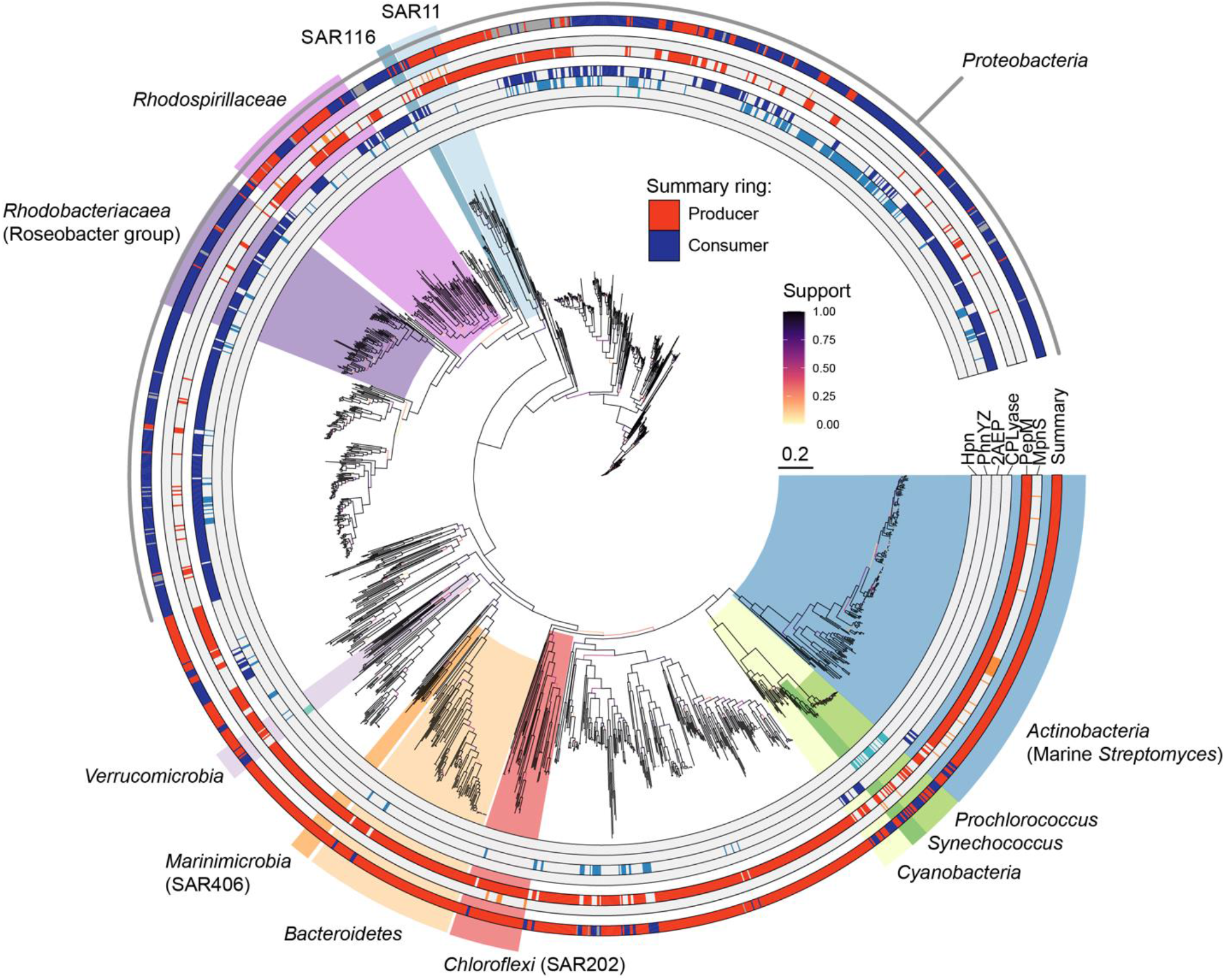
Distribution of phosphonate biosynthesis and utilization pathways in bacterial genomes. Phylogeny is constructed from 120 concatenated, single-copy marker genes from 1890 bacterial genomes from MARMICRODB containing either a phosphonate catabolism or phosphonate biosynthesis pathway. Scale bar is 0.2 amino acid substitution and the tree is unrooted. Monophyletic taxonomic groups with marine representatives are highlighted. The presence of four different phosphonate catabolic pathways (Hpn: Phosphonate catabolism via HpnWXZ, PhnYZ: phosphite or methylphosphonate catabolism via PhnYZ^4^, 2AEP: 2-aminoethylphosphonate catabolism via phosphonoacetate, CPLyase: multisubunit C-P lyase system) is displayed in blue on an inner ring while phosphonate/methylphosphonate biosynthesis pathways (PepM and MpnS) are shown in red on the middle ring. The outer summary ring indicates whether a genome contains at least one catabolic pathway (blue), a phosphonate biosynthesis pathway (red), or both (dark grey, < 0.5% of all genomes). For simplification, the MpnS category also includes the functionally related enzyme HepDI^64^.

### Macromolecular forms of phosphonates

Past studies have linked the biochemical function of phosphonates to their macromolecular form. For example, incorporation of phosphonates into membrane lipids or capsular polysaccharides may protect cells against phospholipase activity or inhibit phage attachment^37–39^. Therefore, we next asked what macromolecular form do phosphonates take in *Prochlorococcus* SB? Phosphonate biosynthesis genes in most *Prochlorococcus* and SAR11 genomes are surrounded with glycosylating enzymes predicted to be involved in the biosynthesis of large extracellular polysaccharide structures, i.e. bacterial capsules (Supplementary Figure S7, Supplementary Table S2). Many marine phytoplankton and bacteria produce extracellular layers of polysaccharides to facilitate aggregation, for defense against predation, and to manufacture biofilms^40^. Phosphonates have been shown to be constituents of the large reservoir of dissolved polysaccharides that accumulate in the surface ocean^5^, and we expected to find phosphonates in *Prochlorococcus* SB to be associated with polysaccharide macromolecules. However, upon fractionation of *Prochlorococcus* organic matter into major biochemical classes, we found that phosphonates were recovered in the protein fraction (Figure 4B) specifically within the methanol/acetone insoluble HMW protein fraction, which includes membrane-associated proteins.

We reconcile these results by proposing that, in *Prochlorococcus* SB, phosphonates are integrated into glycan polymer chains that are then post-translationally attached to HMW proteins, likely membrane-anchored proteins. Indeed, one of the most common post-translational modifications of bacterial proteins is O-linked glycosylation^41^, where glycan polymers are covalently bound to serine or threonine side-chains of larger protein complexes. The phosphonate gene clusters in *Prochlorococcus* SB, SAR11 strain HTCC7217, and SAR11 RS40 genomes are adjacent to glycan assembly enzymes and lipid carriers predicted to be involved in the biosynthesis of capsules (Supplementary Figure S7). However, in some bacteria the glycan building blocks for capsule biosynthesis are also routed towards post translational O-linked glycosylation of membrane-bound lipoproteins^42^. *Prochlorococcus* SB genome contains an O-oligosaccharyltransferase protein directly upstream of the PepM, Ppd, and Pdh cluster. This protein family was first characterized in the attachment of glycans to liposaccharides, but O-oligosaccharyltransferases have more recently been demonstrated to transfer preassembled glycan chains onto protein substrates^43,44^. O-oligosaccharyltransferase genes are also present near PepM in *Pelagibacter* strains HTCC7217 and RS40 (Supplementary Figure S7), and 15% of GORG-tropics SAGs contain PepM plus an O-oligosaccharyltransferase-like domain colocalized within the same 10kbp genome segment. This proportion increases to 50% if we relax the condition that the genes must occur on the same genomic contig. The genomic evidence implies that phosphonates could be a common moiety involved in the post-translational O-linked glycosylation of proteins in the ocean.

### Functional roles of phosphonylated glycoproteins

Why would an oligotrophic-adapted organism like *Prochlorococcus* use scarce P to produce large amounts of phosphonate-containing glycoproteins? Phosphonates are often incorporated into cell-surface structures^37,45^ because they are highly resistant to hydrolysis and can inhibit the activity of some hydrolytic enzymes by mimicking carboxylic acids and phosphate esters^46^. For planktonic bacteria in the ocean, surface-expressed structures provide protection against enzymatic attack, UV-radiation, phages^47^, and protozoan grazers^48^. Modifying cell-surface structures with phosphonates may play a role in reducing vulnerability to grazers and phage, two major drivers of mortality for *Prochlorococcus*^49^. Given that surface expressed proteins are some of the most abundant proteins associated with the cell^50^, the large proportion of cellular P devoted to phosphonates in *Prochlorococcus* SB (Figure 4B) is also consistent with surface protein modification.

Natural selection generally promotes high turnover and low population frequency of genes involved in phage - and predator-interactions^51^ (Supplementary Note S4). Horizontal gene exchange is a key mutational mechanism maintaining genes at low frequencies within microbial populations^51^, and cell surface modification traits are highly enriched in horizontal gene exchange networks^52^. Similarly, we find that patterns of phosphonate biosynthesis gene flow in marine microbial communities do not follow a tree-like pattern and are better explained by horizontal exchange and/or gene loss. Generally, there is a poor correlation between the species tree, inferred from conserved marker genes, and the PepM tree. The topological distance between the species phylogeny and the PepM phylogeny approaches values for random simulated tree sets (Supplementary Table S5). In *Prochlorococcus*, nearly all phosphonate biosynthesis gene cassettes are located within “genomic islands” (Supplementary Figure S8), which contain the majority of laterally transferred genes in *Prochlorococcus*^53,54^. Indeed, there is also a transposase, an important molecular mechanism of lateral gene transfer in bacteria^55^, four genes upstream of PepM in the *Prochlorococcus* SB genome. Taken together, the low frequency of phosphonate producers in bacterial populations and the evidence for horizontal transfer is consistent with a role for phosphonates within cell surface structures – potentially as a defense against phage or grazers.

### Biogeochemical implications: a microbial phosphonate loop within the marine phosphorus cycle

Above 50 meters we expect that chronic, long-term P scarcity drives the cost of phosphonate synthesis to outweigh its fitness benefit for most of the bacterioplankton. This is consistent with the numerical dominance of small, genome-streamlined cells, the low gene frequency of the phosphonate biosynthesis pathway, the low measured concentrations of particulate phosphonates, and the low rates of phosphorus reduction relative to rates of community phosphate uptake in the upper euphotic zone^56,57^. At the nutrient-driven genomic transition zone and near the DCM, organic matter remineralization rates are relatively high and the bacterioplankton community is in close proximity to episodic phosphate inputs from below. Phosphate concentrations begin to rise, and the fitness benefit of phosphonate biosynthesis as a defense against viral lysis and grazing begins to outweigh its cost (Figure 6), sustaining the high frequency of phosphonate producers we found in this depth range. We expect that phosphonates will be most abundant in particulate matter collected near the genomic transition zone, and that higher rates of phosphonate production would lead to higher rates of phosphonate degradation, consistent with the higher C-P lyase gene copy numbers and higher C-P lyase activity measured in microbes inhabiting the deep chlorophyll maximum^7,58^. If phosphonates are used to modify cell-surface glycoproteins as we suggest, then after cell death the labile protein component likely turns over quickly, but the inherently slow degradation of structural polysaccharides would lead to the accumulation of phosphonoglycans as high molecular weight dissolved organic matter^59^ explaining their abundance in marine DOP (Figure 6A). Therefore, a large fraction of dissolved organic P in the ocean may be a direct result of grazer and viral defense mechanisms deployed by a small fraction of marine microbes that allocate a significant amount of their cellular P to phosphonate production.

**Figure 6:**
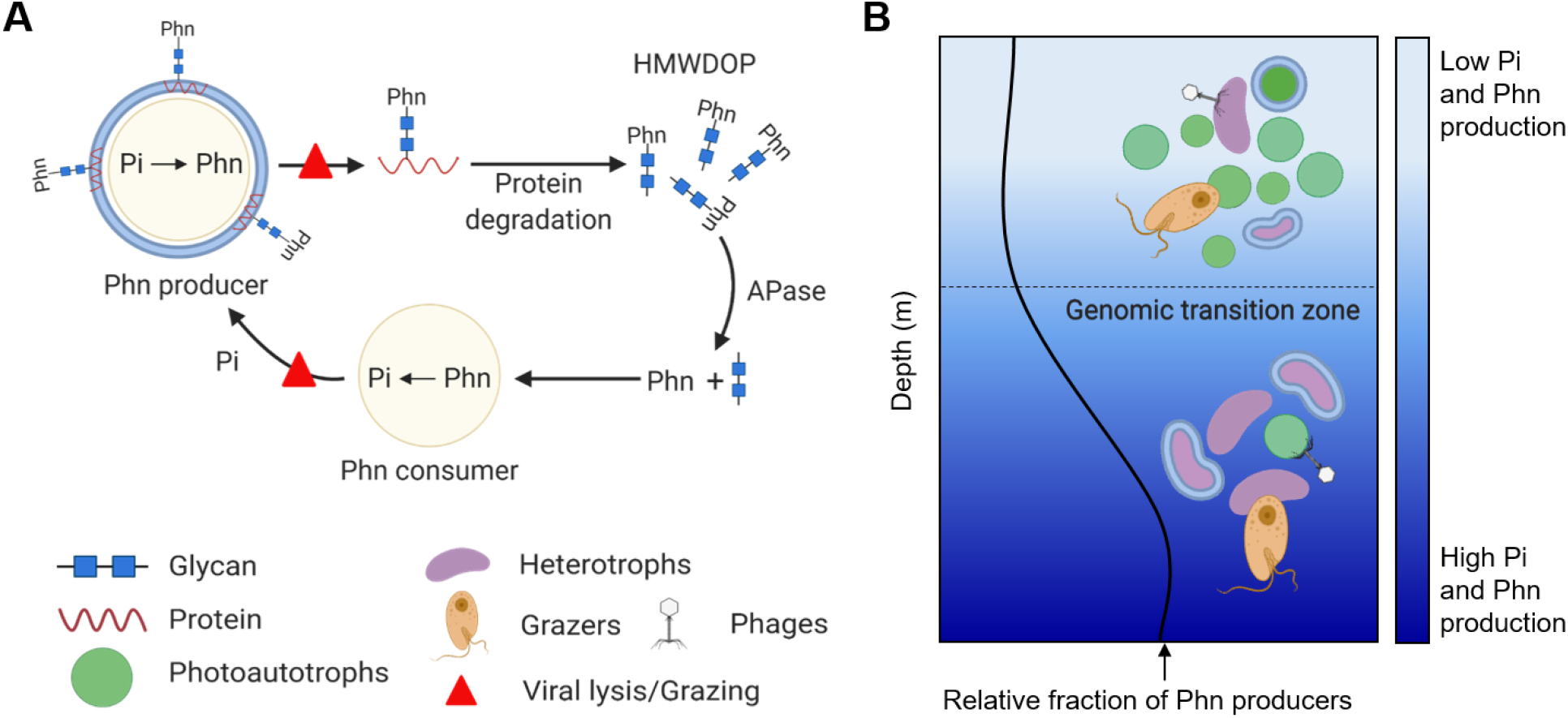
Microbial cycling of phosphonates in the upper ocean. **A)** Microbes with PepM produce cell surface-layer phosphonoglycoproteins to reduce mortality from grazing or viral infection. Upon death of the cell phosphonoglycoproteins are released into seawater where heterotrophic microbes quickly remineralize proteins leaving phosphonoglycans to accumulate as high molecular weight dissolved organic phosphorus (HMWDOP). Phosphonates are hydrolyzed from glycans by alkaline phosphatases (APase) family enzymes and further hydrolyzed into inorganic phosphate (Pi) by C-P lyase or other hydrolytic pathways. Recycled Pi can then be used to produce new phosphonates. **B)** In the surface ocean, Pi is scarce and often limiting. Producing phosphonate as a mortality defense may be costly in terms of resource allocation. Therefore, the relative fraction of phosphonate producers in the microbial community is low. As nutrient availability increases through the genomic transition zone^27^ the benefit/cost ratio of phosphonate production increases and phosphonate producers are relatively more abundant.

## Acknowledgments

We thank A. Coe, K. Dooley and C. Bliem for their invaluable help with *Prochlorococcus* cultivation; E. M. Grabowski, K. M. Bjorkman and D. M. Karl for their measurements of particulate carbon and phosphorus; C. G. Johnson for his assistance with the NMR. This work was supported in part by grants from the National Science Foundation (OCE-1153588, and DBI-0424599 to S.W.C.; OCE-1335810 and OIA-1826734 to R.S. and OCE-1634080 to D.J.R), the Gordon and Betty Moore Foundation (#6000 to D.J.R) and the Simons Foundation (Life Sciences Project Award IDs 337262, 647135, S.W.C.; 510023, R.S.; SCOPE Award ID 329108, S.W.C and D.J.R.). This paper is a contribution from the Simons Collaboration on Ocean Processes and Ecology (SCOPE).

## Authors contributions

M. Acker grew the cultures, performed the macromolecular fractionation and protein extraction, acquired and interpreted the NMR spectra. S. L. Hogle performed all bioinformatics and biostatistics. T. Hackl assisted with genomic island prediction. P. M. Berube identified *Prochlorococcus* SB as a potential phosphonate producer and provided cellular biomass to D.J. Repeta for preliminary experiments. R. Stepanauskas provided early access to the GORG-Tropics database. S. W. Chisholm supported the culture work by providing laboratory space, training and advice. D. J. Repeta conceptualized the study, helped with data interpretation, and provided general guidance on the execution of experiments. M. Acker, S. L. Hogle and D.J. Repeta drafted the manuscript with assistance from all the co-authors.

## Methods

### Genomic data sources

We used two collections of genomes for the comparative genomics analysis: MARMICRODB^36^ (https://zenodo.org/record/3520509) and the Global Ocean Reference Genomes (GORG) Tropics dataset^15^. MARMICRODB contains over 18000 archaeal, bacterial, eukaryotic, and viral genomes from predominantly the marine environment, but also terrestrial and host-associated systems. GORG Tropics consists of approximately 13000 single-cell genomes sequenced from 28 samples across the tropical surface ocean. Roughly 6000 genomes from GORG Tropics come from a single sample (GORG BATS) from the Sargasso Sea. For quantitative analysis of the complete GORG dataset we randomly subsampled GORG BATS (Sample SWC-09) genomes to the median genomes per sample (N=241) from the remainder of GORG Tropics. We also used 195 surface and Deep Chlorophyll Maximum metagenomes from Tara Oceans project^18^ and 480 metagenomes from bioGEOTRACES, HOT, and BATS^17^. Genome and metagenome quality control and exclusion criteria were performed as described earlier^36^.

### Homology searches

Phosphonate biosynthesis and catabolism proteins were identified by homology to a collection of hidden Markov models using HMMERv3.1b2 (http://hmmer.org/) and the trusted cutoffs of each individual model. PepM sequences were initially identified using PF13714 (PEP_mutase) and then aligned to the Tigrfam^60^ Hidden Markov Model (HMM) TIGR02320 using hmmalign. Authentic PepM sequences were identified by strict presence of conserved active site motif “EDK(X)5NS” or if they lacked at most two active site residues but were located adjacent (within 5000 nucleotides upstream/downstream) to a gene coding for Ppd (TIGR03297) or MpnS^14,61–63^. The PF13714 model identified many sequences lacking the PepM active site motif, and with significant similarity to the related isocitrate lyase superfamily PF00463. Sequences passing the PF13714 bitscore cutoffs, but lacking the “EDK(X)5NS” motif were classified as members of the isocitrate lyase superfamily. Sequences from authentic MpnS and the related HepDI/HepDII proteins were identified using custom HMMs built from alignments of curated sequences identified as containing essential catalytic residues^64^. These alignments were constructed using MAFFT v7.273^65^ in E-INS-i iterative refinement mode. The alignment was trimmed to peptide positions 317-792 to isolate the core catalytic residues and used to construct HMMs. MpnS, HepDI, and HepDII sequences in MARMICRODB and GORG-tropics were identified using the resulting HMMs (evalue < 0.05), and the resulting sequences were aligned using MAFFT. The alignments were manually inspected to remove sequences lacking the essential catalytic residues and were iteratively realigned after poorly-aligned sequences were removed.

For phosphonate catabolism proteins, gene neighborhoods were inspected to confirm the presence of multiple co-occurring genes from the PhnYZ^4^, CP lyase, 2-aminoethylphosphonate, and phosphonoacetaldehyde phosphonatase catabolism pathways. Specifically, valid C-P lyase clusters required at least five other genes from phnC, phnD, phnE, phnF, phnG, phnH, phnI, phnJ, phnK, phnL, phnM, phnN, phnO, phnP to occur within uninterrupted (ie no scaffold/contig breaks) 16000 nucleotide sliding windows. Valid phosphite clusters required two of ptxD, phnY, and phnZ. Valid 2-aminoethylphosphonate clusters required co-occurring phnX and phnW or phnW, phnA, and phnY. Valid phosphonoacetaldehyde phosphonatase clusters required phnZ, hpnW, and hpnZ. All catabolic pathways other than C-P lyase required genes to co-occur within uninterrupted 10000 nucleotide sliding windows.

We also identified two sets of ten highly conserved, single-copy families from *Prochlorococcus* and SAR11 lineages for use in normalization with metagenome profiling. These gene families had the ten lowest dN/dS ratios of the entire pangenome, that is the highest purifying/stabilizing selection, for *Prochlorococcus* or SAR11, and should thus be highly specific and sensitive for metagenomic quantification. *Prochlorococcus* and SAR11 pangenomes were characterized and defined using PanX^66^. To profile metagenomes for total bacterioplankton, we used fetchMG from the mOTUs2 tool^67^. Briefly, we identified ten COG families (COG0012, COG0016, COG0018, COG0172, COG0215, COG0495, COG0525, COG0533, COG0541, COG0552) from all bacterial and archaeal genomes in MARMICRODB. To reduce redundancy, we clustered protein sequences from each COG at 90% sequence identity using MMseqs2 v4e23d^68^and used the 90% clustered sequence representatives in metagenome search databases.

The HMM protein sequence profiles, results from these searches, and descriptions of the sequence families are available from https://github.com/slhogle/phosphonates.

### Multiple sequence alignments, phylogenetic inference, and topological comparison

Authentic PepM sequences containing the “EDK(X)5NS” motif were aligned to TIGR02320 using hmmalign and trimmed using trimAl v1.4.rev22^69^ with the automated-gappyout option, and alignments were inspected manually to ensure the veracity of the “EDK(X)5NS” motif. The multi-phylum genome phylogenies were created from genome assemblies with confirmed phosphonate biosynthesis pathway or at least one confirmed phosphonate catabolic pathway. We used the GTDB-Tk v1.3.0 pipeline^70^ with default settings and the GTDB R05-RS95 database^71^ to identify conserved proteins (120 bacterial proteins/122 archaeal proteins) and generate concatenated multi-protein alignments. We filtered alignment columns using the bacterial and archaeal alignment masks from (http://gtdb.ecogenomic.org/downloads). We then removed columns represented by fewer than 50% of all taxa and/or columns with no single amino acid residue occurring at a frequency greater than 25%. We further trimmed the alignments using trimAl with the automated - gappyout option to trim columns based on their gap distribution. The multi-phylum genome phylogenies and the PepM phylogeny were inferred using FastTree v2.1.10^72^ under the GAMMA model of rate heterogeneity and the WAG+ substitution model^73^. Support values were determined using 100 non-parametric bootstrap replicates. Both the PepM tree and genome tree were left unrooted. Phylogenies and associated data were visualized using ggtree^74^. Detailed phylogenies of *Prochlorococcus* and *Pelagibacterales*/SAR11 in the supplementary materials were constructed as described earlier^36^.

Comparisons between the topology of the PepM tree and the genome phylogeny were performed using ETE3 v3.1.1^75^. We pruned both the PepM and genome phylogenies so that they contained the same number taxa and no duplication events in the PepM tree, which resulted in 999 leaves for bacteria and 16 leaves for archaea. We then compared the topologies of the PepM tree, the genome tree, and ten simulated random trees using the Robinson-Foulds symmetric distance^76^ and the fraction of edge similarity. For Robinson-Foulds distances, edges were treated as unpolarized splits rather than “true” clades because all trees were compared as unrooted. The Robinson-Foulds distance simply counts the number of branch partitions (nodes) that appear in one tree but not the other. Therefore, the maximum possible Robinson-Foulds distances of an n-taxa unrooted tree is 2(n-3). To compute a normalized distance, we simply divided the observed Robinson-Foulds distance by the maximum distance of the two-way comparison between trees. Thus, the normalized Robinson-Foulds distance is a value from 0 to 1, which can be interpreted as the fraction of nodes/splits missing in the query tree compared with the reference tree.

### Gene enrichment analysis

We annotated the 16 *Prochlorococcus* and 22 SAR11/*Pelagibacterales* genomes containing verified PepM sequences with eggNOG 4.5.1^77^ using eggNOG-Mapper v1.0.3-3-g3e22728^78^. We then collected the resulting KEGG orthology annotations^79^ from all genes and tested for enrichment of modules and pathways in the KEGG hierarchy in genes within 10000 nucleotides upstream/downstream of the PepM sequence start and stop coordinates respectively. Significant enrichment of KEGG categories were determined using the hypergeometric test^80^ implemented in clusterProfiler v3.8^81^. We excluded genomes from the analysis that had contig/scaffold breaks within 10000 nucleotides upstream/downstream of the PepM sequence.

### Identification of PepM sequences within Prochlorococcus genomic islands

We predicted genomic islands using Hidden Markov Models trained from conserved gene synteny patterns in closed Prochlorococcus genomes. Briefly, we assume genomic islands to be contiguous stretches of DNA enriched in flexible genes families (i.e. genes found in a subset of all genomes). We then used previously described genomic islands in six Prochlorococcus genomes^54,82,83^ to define core, flexible, and inconclusive gene “states” in four distinct hidden markov models, each with two hidden states (island or non-island). We then used empirical gene family frequencies as a proxy for the core/flexible/inconclusive state of each gene and the gene order on each scaffold as input to the Viterbi algorithm for predicting the hidden island state of each gene in all other *Prochlorococcus* genomes.

### Estimation of PepM prevalence in *Prochlorococcus* and SAR11 genomes

Because many of the *Prochlorococcus* and SAR11 genomes analyzed here are incomplete (SAGs or MAGs), we attempted to estimate the ‘true’ proportion of genomes with PepM while correcting for genome incompleteness. Briefly, we used the estimated genome completeness from checkM v1.0.11^84^, which is based on the presence of core marker genes, to estimate the number of missing bases per taxonomic group (*Prochlorococcus* or SAR11) and then use this scale the relative abundance of potential phosphonate producers per group. We estimated the corrected prevalence of phosphonate producers as:

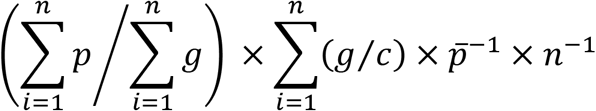

where for each clade, *p* is the length of the phosphonate biosynthesis/degradation operon in base pairs, *g* is the length of the genome assembly in base pairs, *c* is the completeness estimate from CheckM, p is the average length of all phosphonate biosynthesis/degradation operons from each clade, and *n* is the total number of assemblies from each clade.

### Metagenome read classification

We created Diamond v0.9.22.123^85^ search databases for each of the three ten-gene marker gene sets identified for *Prochlorococcus*, SAR11, and all bacterioplankton using reference genomes from MARMICRODB. We also created Diamond databases for only *Prochlorococcus* PepM, only SAR11 PepM, and all bacterial and archaeal PepM sequences identified with the “EDK(X)5NS” catalytic motif from MARMICRODB only. We identified the marker gene sets using a two-tiered search strategy using Diamond in default fast mode. We first searched the metagenomes against reduced marker sets clustered at 70% amino acid identity by MMseqs2. We pulled all reads with hits to these reduced marker sets, then searched the reads against the entire MARMICRODB, and finally retained read mappings with a best scoring match to a MARMICRODB protein from the original marker family. We searched metagenomic reads against the PepM databases using Diamond (mode --more-sensitive) and score cutoffs of 55% amino acid identity and an E-value of 1e-5. We determined these cutoffs empirically to be those that produced the highest F Score (harmonic mean of precision and recall) from mock metagenomes simulated from GORG-tropics (Supplementary Note S5, Supplementary Figure S9). Since the E-value is a function of database size it is important to note that the significance of this cutoff is specific to the reference databases here. In the case of ambiguous alignments (i.e., identical best alignment scores to sequences from different taxonomic groups) we classified reads using a probability-weighted random sampling of all taxonomic groups matching the best hits. We derived the probability distributions for random samplings from the taxonomic classification uniquely mapped PepM reads.

To normalize PepM reads across metagenomes we calculated the length normalized counts (RPKM) of the 10 single copy marker genes from *Prochlorococcus*, SAR11, and all bacteria and archaea in each metagenome as reads per kilobase marker and then estimated the number of ‘genome equivalents’ in each metagenome as the median of the 10 marker families. We also calculated the length normalized abundance of PepM from *Prochlorococcus*, SAR11, and all bacteria and archaea as reads per kilobase from each metagenome. We then divided the length normalized abundance of PepM by the estimated number of genome equivalents for each taxonomic group to estimate the fraction of genomes with phosphonate biosynthesis potential. To ensure robust estimates of normalized abundance we excluded samples where the median marker gene coverage was less than 100X for all taxonomic groups.

### Biotic and abiotic data associated with metagenomes

The biotic and abiotic variables used in this study were obtained and preprocessed as described in detail here: https://doi.org/10.5281/zenodo.3689249 and here: https://doi.org/10.5281/zenodo.3786232. We obtained phosphate concentrations from the GEOTRACES Intermediate Data Product IDP2017 version 3^22^, specifically from sections GA02^86,87^, GA03, GA10^88^, and GP13. We obtained dissolved phosphate concentrations from the Tara Oceans project^89^ (https://doi.pangaea.de/10.1594/PANGAEA.875579). Modeled climatological dissolved organic phosphorus and other variables were obtained from the MIT Darwin model (v0.1_llc90, http://darwinproject.mit.edu/) from the Simons Collaborative Marine Atlas project (CMAP) https://simonscmap.com/ using pycmap v0.1.2 (https://doi.org/10.5281/zenodo.3561147). Taxonomic profiles derived from metagenomic reads were generated as described earlier^36^ using the MARMICRODB database. As a first order estimate of *Prochlorococcus* and SAR11 ecotype relative abundance we divided the number of reads mapping to each ecotype by the number of reads mapping in total the *Prochlorococcus* genus or the family *Pelagibacterales* for SAR11. Some proportion of reads mapping to highly conserved (core) regions can only be reliably classified at the genus or family level thus our ecotype relative abundance estimates using total read counts are underestimates.

### Random Forest Regression

We trained one random forest model (n_trees_= 1000) for each taxonomic group; *Prochlorococcus*, SAR11, and bacteria plus archaea. Each model was trained using up to 44 abiotic/biotic variables including trace metal and macronutrient data from GEOTRACES and Tara Oceans, modeled climatological means from the MIT Darwin model (http://darwinproject.mit.edu/), and ecotype relative abundances. For the outcome variable (normalized PepM relative abundance) we used a modified splitting rule for tree construction that maximized the log-likelihood of the beta distribution on the interval [0,1]^90^. Although our relative abundance measure is not theoretically restricted to the unit interval (values greater than one could exist, e.g. if the PepM copies per genome greatly exceeded one), in practice PepM relative abundance was always bounded [0,1] in our datasets. We performed random forest regression using nested ten-fold cross-validation to prevent data leakage to the validation phase^91^. We reserved 20% of the data for estimating final model performance. The remaining 80% of training data was split into 10 resampled partitions, each with analysis and assessment partitions, to tune and estimate the performance of preprocessing, supervised feature selection, hyperparameter tuning steps. We tuned hyperparameters (mtry, min.node.size), by maximizing the coefficient of determination from correlation (model R^2^) and minimizing the root mean square error (RMSE). We used CAR Scores^92^ for recursive feature elimination to retain only the top 50th percentile of informative variables. We included the feature elimination step to reduce the computation costs and runtime of the feature importance step (see below). Random forest regression was implemented with the package Ranger^25^ and the Beta Forest algorithm^90^. We determined predictor variable rankings on the final model from the cross validation step using the Boruta heuristic^93^. This step allowed us to identify all predictor variables that consistently performed better than chance and to compare the importance of each variable to a reference importance level, i.e. random data.

### Beta-Binomial Regression and Generalized Additive Models

We used the R package corncob^26^ and modeled PepM relative abundance directly from PepM read counts and the median read counts to marker gene sets as “successes/total” which is appropriate for the beta-binomial probability distribution. We used the log-odds link function for both relative abundance and the overdispersion parameter. For each taxonomic group (SAR11/*Prochlorococcus*/bacteria and archaea) we modeled only the top five most important biotic/abiotic variables identified in the Random Forest regression and variable importance steps. We did not include additional model terms because multiple collinearities between covariates and the additional model complexity prevented model convergence in most cases. We estimated the probability for each biotic/abiotic covariate being informative to the overall model by using bootstrapped likelihood ratio tests (N=1000). We estimated 95% confidence intervals for model coefficients and standard errors using 1000 random draws from the beta-binomial distribution.

We estimated seasonal effects in time-series metagenomes using Generalized Additive Mixed Models and Linear Mixed-Effect Models implemented through the mgcv v1.8-26^94^ nlme v3.1-148 libraries in R v3.6.2. To decompose any potential seasonal effects, we fit a cyclic spline term to a variable for the day of the year, which we use as a proxy for season, and we fit a global trend term to the cumulative time since sampling onset. We considered a seasonal effect present if the model term for “day of the year” was statistically significant (p < 0.05).

### *Prochlorococcus* cultures under P-replete and P-deficient conditions

To investigate the linkage of PepM with phosphonate production, we grew two HLII strains of *Prochlorococcus*: *Prochlorococcus* SB, which has PepM, and the closely related *Prochlororoccus* MIT9301, which lacks the PepM gene sequence. Both strains were grown axenically, under constant light (30 μmol quanta m^−2^ s^−1^) in artificial seawater medium AMP1 prepared has described before^95^, but using 3.75 μM TAPS as a buffer instead of 1mM HEPES. After growing *Prochlorococcus* SB in the regular medium i.e with a phosphate concentration of 50 μM (N/P = 16/1) to assess whether or not this strain could produce phosphonate, we decreased the phosphate concentration to 2.28 μM (N/P = 350/1). Although the phosphate concentration was lower, *Prochlorococcus* SB cells were not P-limited in exponential phase growth. However, stationary phase was induced by P-starvation, as demonstrated by the initiation of further growth after adding phosphate to the culture (Figure 4A inset). For all experiments, biological duplicates were grown to ensure reproducibility. Cultures axenicity was assessed by flowcytometry and by confirming a lack of turbidity for at least 30 days after inoculation with three test broths: ProAC^96^, MPTB^97^ and ProMM (Pro99 medium^95^ supplemented with 1xVa vitamin mix^98^ and 0.05% w/v each of pyruvate, acetate, lactate and glycerol. ProMM is the 100% seawater based version of the PLAG medium^96^. All the glassware and polycarbonate bottles (1 L for the blank, 2 L and 20 L for the cultures) were cleaned by soaking overnight in 2% detergent (micro), rinsed 6 times with deionized water, soaked overnight in 1 M HCl and rinsed 6 times with ultra-high purity water.

### Cell harvest and treatment

*Prochlorococcus* SB cultures grown in high N/P medium were harvested twice: ~6 L were harvested during exponential growth and the remainder (14 L) harvested two days after the onset of stationary phase growth. To ensure that stationary phase cells were limited by phosphorus, 25 mL of culture was amended with phosphate. Fluorescence increased in the phosphate-amended cultures, reaching levels regularly observed in *Prochlorococcus* HLII cultures (Figure 4A inset). Cells were separated from the growth medium by centrifugation (15,970 rcf for 30 minutes at 4°C) and the growth medium was saved for other analyses. Cell pellets were transferred into 50 mL falcon tubes, suspended in Turk Island mix^95^ to rinse the cells of external nutrients and centrifuged (6,523 rcf for 15 minutes at 15°C). This time, we discarded the supernatant and repeated the operation two more times. After, we flash froze the cell pellets in liquid nitrogen and stored them at −20°C until NMR analyses. We also measured *Prochlorococcus* cell abundance by flow cytometry. Samples were prepared and processed as previously described^99,100^ a d run on an Guava 12HT flow cytometer (Luminex Corp., Austin, TX, USA). Cells were excited with a blue 488 nm laser analyzed for chlorophyll fluorescence (692/40nm), SYBR Green I stained DNA fluorescence content (530/40nm), and size (forward scatter). Samples were ran stained with 1x SYBR Green I (Invitrogen, Grand Island, NY) and unstained then incubated for 60 min in the dark prior to running. All flow cytometry files were analyzed using Guavacyte.

### Nuclear Magnetic Resonance

NMR spectra were acquired at 25°C on a 400 MHz Bruker AVANCE DPX spectrometer using a 5mm inverse broadband probe and running TOPSPIN 1.3. ^31^P shifts are reported relative to external 85% phosphoric acid at 0 ppm. For the proton-decoupled ^31^P-NMR spectra, we used ‘zgdc30’ with WALTZ16 decoupling and sweep width of 80 ppm, a 3 seconds relaxation delay, 100K scans and 20Hz line broadening. *Prochlorococcus* SB and MIT9301 whole cells were packed into a 5mm BMS tube (Shigemi Inc.) with magnetic susceptibility of the glass inserts matching D_2_O.

^31^P-NMR spectra of *Prochlorococcus* SB protein fraction was acquired at 25°C on a 400 MHz Bruker Ascend 400 equipped with a Sample CASE. The ^31^P-NMR spectra were acquired using the program ‘zgpg30’ with a sweep width of 80 ppm, a relaxation delay of 2 seconds, a 15Hz line broadening and for 13K scans. ^31^P chemical shifts are reported relative to external phosphoric acid at 0 ppm.

### Elemental composition of *Prochlorococcus* SB and MIT 9301

Elemental C/N/P ratios were measured at the University of Hawai’i nutrient facility according to the protocols employed by the Hawaii Ocean Time series program (http://hahana.soest.hawaii.edu/hot/protocols/protocols.html#). Briefly, cell pellets from ~ 900 mL of culture were transferred to combusted glass vials, dried, and powdered. C and N were measured on subsamples using a PE-2400 Carbon/Nitrogen analyzer calibrated with acetanilide standards. Cellular P was measured by the molybdenum blue method^101,102^ after first combusting cell pellets at 450°C for 3h, and dissolving the residue in 10 mL of 0.5 M HCl.

### Separation of cellular macromolecular classes

To fractionate *Prochlorococcus* organic matter into different classes of major biochemicals, we followed the protocols of Karl et al.^103^. Cells from 3 L of culture were centrifuged and the isolated cell pellet extracted with 5 mL of cold 5% trichloroacetic acid (TCA) for 1h. The mixture was centrifuged (12,100 rcf for 30 minutes at 4°C), the supernate decanted, and the TCA insoluble material washed twice with 5% cold TCA. The TCA fractions were combined, evaporated to dryness. Lipids were recovered from the TCA insoluble material by extraction (3x, room temperature, 20 minutes each) with 95% ethanol (5 mL). Residual ethanol was evaporated, and RNA within the dry TCA/95% ethanol insoluble material hydrolyzed at 37°C for 1 hour with 2.5 mL of 1 M NaOH. The hydrolysis was quenched by immersing the sample tube in an ice bath for 15 minutes, after which the sample was acidified to pH 1 by adding 2.5 mL 1 M HCl and 0.5 mL of 50% TCA. The mixture was allowed to sit for 15 min to precipitate proteins and DNA, then centrifuged for 30 minutes at 12,100 rcf. After collecting the supernatant (containing RNA), we rinsed and centrifuged the pellet 2x with 5 mL of 5% TCA and 2x with 5 mL of 95% ethanol. To hydrolyze DNA, we added 5 mL of 5% TCA to the insoluble material and immersed the tube in boiling water for 30 minutes. After centrifuging the tube and collecting the DNA-containing supernatant, we rinsed and centrifuged the pellet 2x with 5 mL of ice-cold 5% TCA and 2x with ice-cold 95% ethanol as before. Finally, we extracted the protein from the remaining cell pellet with 5 mL of 1 M NaOH (37°C,18 h). A small amount of insoluble debris remained after the protein extraction. This was removed by centrifugation (12,100 rcf; 30 minutes) and followed by syringe filtration of the supernatant.

### Protein extraction and precipitation

Peptides and denatured “soluble” proteins were fractionated from native HMW “insoluble” proteins using the protocol described by Hutchins et al.^104^. The cell pellet of 0.5 L of culture was lysed (15 minutes, RT) with 1 mL of 1% SDS extraction buffer (1% SDS, 0.1 M Tris/HCl pH 7.5, 10 mM EDTA) then heated at 95°C for 10 minutes. Samples were allowed to cool to RT and were then agitated at 350 rpm for 1 hour. The resulting suspension was centrifuged (20 minutes, 14,100 rcf), and the supernatant decanted. After transferring the supernatant containing the proteins, we concentrated the proteins from the supernate by membrane centrifugation using 5 K molecular weight cutoff Vivaspin units of 6 mL (Sartorius Stedim, Goettingen, Germany). The retentate (~300 μL) was recovered, and 1 mL of cold 50/50 methanol/acetone solution (acidified with HCl to a final concentration of 0.5 mM) added. After sitting for 3 days at –20°C, insoluble proteins were pelleted by centrifuge (14,100 rcf for 30 minutes at 4°C) and the supernatant decanted.

### Dataset/code availability

The datasets and computer code supporting the findings in this study are available from: https://github.com/slhogle/phosphonates.

The entire MARMICRODB dataset including a comprehensive description, raw protein fasta files, Kaiju v1.6.0^105^ formatted databases, scripts and instructions for how to use the resource is available from https://doi.org/10.5281/zenodo.3520509.

GEOTRACES chemical data was processed and matched to metagenome samples using code/methods available from https://doi.org/10.5281/zenodo.3689249.

Tara Oceans chemical and hydrographic data was processed and matched to metagenome samples using code/methods available from https://doi.org/10.5281/zenodo.3786232.

The list of *Prochlorococcus* core PFAM and TIGRFAM families, a compiled HMMERv3 hidden Markov model database, and a CheckM formatted marker list file is available from https://doi.org/10.5281/zenodo.3719132.

