## Supplementary Material for "Phosphonate production by marine microbes: exploring new sources and potential function"

**Supplementary Notes**

**Supplementary Note S1: Taxonomic breadth of marine phosphonate producers and consumers**

Phosphonate producers in the GORG dataset are more diverse than consumers. We identified phosphonate producers in 30 unique families and 11 different phyla (*n* domain=2; *n* phylum=11; *n* class=12; *n* order=25; *n* family=30; *n* genus=40; total genomes=6720). Producers are from diverse groups including the Alphaproteobacteria, Gammaproteobacteria, Flavobacteriales, Bacteroidia, Actinobacteria*,* Archaea*,* and Cyanobacteria, and also uncultured groups like SAR202 and the candidate phyla Marinamargulisbacteria and Marinisomatia (Figure 2). While for phosphonate consumers, we identified 13 unique families and 5 different phyla (*n* domain=1; *n* phylum=5; *n* class=5; *n* order=10; *n* family=13; *n* genus=17; total genomes=6720). However, the distribution of taxonomic ranks for phosphonate producers was less even than for consumers. Over 60% of phosphonate producers were assigned to the SAR11 group, mostly clade Ia.3, while the remaining 40% were evenly distributed amongst low frequency families/orders. The ZD0417 (6%) clade, the OM1 clade (4%) and High-light *Prochlorococcus* (4%) were the next most abundant producer groups. In contrast, 30% of phosphonate consumers are from the Aegean-169 clade (including *Alphaproteobacterium* HIMB59)[1], 21% from the *Rhodobacteraceae*, 18% from SAR11 clade Ia.3, 8% from SAR11 clade IV, and 12% were High-light *Prochlorococcus*.

In MARMICRODB we found phosphonate producers in most of the same taxonomic groups as the GORG database. Unlike GORG, MARMICRODB also includes many genomes isolated from the deep ocean (> 500 meters) and from marine sediments. We found abundant production potential in many deep ocean planktonic groups such as Chloroflexi (including the SAR202 clade) and Marinimicrobia (SAR406). We also found PepM and the Ppd expressed in 24/186 marine transcriptomes from the MMETSP[2] mostly from dinoflagellates as well as in the abundant and cosmopolitan marine Prasinophyte, *Ostreococcus*. Relative abundances estimated from MARMICRODB alone should be viewed with caution since MARMICRODB has a biased composition. However, we note that fewer than 1.5% of the archaeal genomes in MARMICRODB were phosphonate producers (n=18/1256) which is in strong agreement with the distributions in GORG-tropics. Like GORG-tropics, no archaeal genomes from MARMICRODB contained phosphonate catabolism pathways.

**Supplementary Note S2: Seasonal dependence of phosphonate producers at BATS**

We examined the abundance of phosphonate producers in two long-standing ocean time-series: The Bermuda Atlantic Time-series Study in the North Atlantic Ocean (Sargasso Sea) and the Hawaii Ocean Time-series near Hawaii in the North Pacific Ocean. Both study sites are in oligotrophic ocean gyres and have comparable biological productivity and carbon export[3]. *Prochlorococcus* and SAR11 are the most abundant planktonic cells in the upper 300 meters at both sites[4]. However, the two study sites have different seasonal patterns. BATS experience strong convective mixing during the winter months, and HOT is more stably stratified, although HOT also displays seasonal productivity cycles[5]. The two study sites also differ in their biogeochemical inventories. Inorganic phosphate concentrations are lower at BATS than HOT which is likely due to the effect of iron supply on dinitrogen fixation rates in each ocean basin[6]. Indeed, atmospheric iron fluxes are higher at BATS compared with HOT[7,8]. Differences in dissolved phosphorus inventories between HOT and BATS have been previously linked to gene content variation between *Prochlorococcus* populations. *Prochlorococcus* populations in the Sargasso sea are more likely to have additional P acquisition genes such as alkaline phosphatase[9,10].

We detected phosphonate producers at both HOT and BATS at similar abundances as in the rest of the global ocean. For SAR11 and total bacterioplankton these abundances generally increased with depth and were highest in the deep mixed layer below the subsurface deep chlorophyll maximum layer (> 200 meters). *Prochlorococcus* phosphonate producers were at highest abundance in the surface layer (approximately 10 meters) at HOT and BATS and decreased in abundance with depth. At HOT approximately a constant 6-9% of *Prochlorococcus* genomes, 10% of SAR11 genomes, and 10% of total bacterioplankton genomes had a phosphonate production gene cluster in the surface layer. At BATS we detected a significant seasonal effect on the abundance of all phosphonate producers in the surface layer that temporally corresponds with the recurring deepening of the mixed layer during winter months (Supplementary Figure S4C). For example, *Prochlorococcus* phosphonate producers are most abundant from February to May then decline to negligible levels in the summer at the onset of peak water column restratification. The seasonality of SAR11 producers and the total community producers was tightly coupled, suggesting that peak producer abundance occurs during periods with the deepest mixed layer (November through the following March). This pattern follows the seasonal succession of ecotypes at BATS where SAR11 clades IIa and IIb are most abundant in the upper 200 meters during the winter[11]. SAR11 clades IIa and IIb were also the most abundant producers in GORG-tropics and GORG-BATS after the surface clade Ia, and SAR11 IIb abundance showed a strong positive correlation with phosphonate producers in the global dataset.

**Supplementary Note S3: Biotic and abiotic covariates shaping the global distribution of phosphonate producers**

For *Prochlorococcus* producers we find that a small, but significant, amount of variation is explained by inorganic phosphate concentrations, and the modeled climatological mean of dissolved organic phosphorus (Supplementary Figure S3 and S5). The models suggest that on average *Prochlorococcus* phosphonate producers are most abundant where inorganic phosphate and dissolved organic phosphorus concentrations are highest. For example, the

North Atlantic and Mediterranean had the lowest median abundance of *Prochlorococcus* producers. In the deeply sequenced N. Atlantic sample GORG-BATS we detected no *Prochlorococcus* phosphonate producer genomes and significantly more *Prochlorococcus* phosphonate consumer genomes than in the rest of the global samples. This pattern mirrors the overall depletion of inorganic phosphate relative to other nutrients in the N. Atlantic[6]. We interpret this as reflecting potential selection against the phosphonate biosynthesis trait in *Prochlorococcus* populations from consistently P-limited regions, such as the tropical N. Atlantic, due to the additional phosphorus cost of phosphonate biosynthesis. The largest proportion of variation for *Prochlorococcus* phosphonate producers is explained by the total abundance of *Prochlorococcus.* That is, *Prochlorococcus* phosphonate producers are more abundant when *Prochlorococcus* constitutes a smaller fraction of the total microbial community. It is unclear what ecological mechanisms underlie this pattern, and it may simply be a consequence of the heteroskedasticity of compositional data, i.e. higher variances as the denominator in the ratio of (PepM/Core genes) approaches zero.

The relative abundance of SAR11 phosphonate producers and the bacterioplankton producers were strongly positively correlated, consistent with SAR11 being the dominant phosphonate producer in GORG-tropics and GORG-BATS. Thus, the relative abundances of SAR11 producers and total producers were shaped by many of the same factors. A small but significant amount of variation in SAR11 phosphonate producers was explained by increasing climatological mean nitrate concentrations and dissolved aluminum concentrations. This reflects that the highest average abundance of SAR11 producers occurred in the North Atlantic and Mediterranean Sea, regions which have higher than average dissolved aluminum concentrations[12] and climatological mean nitrate concentrations in the upper 100 meters. Generally, we find that the greatest amount of phosphonate producer variation is explained by multiple depth-dependent biotic factors implying that increasing depth is a unifying master parameter (Figure 2, Supplementary Figure S2, S3 and S5, main text). Total bacterial/archaeal and SAR11 phosphonate producers increase alongside depth-associated biotic factors including low light *Prochlorococcus* and the abundance of mesopelagic groups like Archaea[13] and SAR11 IIb[14]. Phosphonate producers are negatively associated with known surface clades like SARII IV and V and abiotic factors like dissolved manganese concentrations (manganese being a surface enriched element[15]). These modeling results offer a prediction that bulk phosphonate concentration is greatest in the lower reaches of the euphotic zone and upper reaches of the mesopelagic. Future chemical studies targeting the 100-300 meters depth range will hopefully provide greater understanding of phosphonate cycling in the upper ocean.

### Supplementary Note S4: Evidence for phosphonylated surface layer structures by ***Prochlorococcus*** SB

The phosphonate biosynthesis gene cluster contains genes predicted to be involved in cell wall/membrane/envelope biogenesis and extracellular polysaccharide biosynthetic processes (Supplementary Table S2), suggesting that *Prochlorococcus* SB may produce extracellular saccharides modified with phosphonate groups. We also found that genes coding for glycosylation reactions were statistically more abundant within ± 5 Kbp of the phosphonate biosynthesis cluster than in the remainder of the complete genomes from *Prochlorococcu*s SB and *Pelagibacter* sp. HTCC7217 and RS40 (Supplementary Table S2). Many of these genes encode glycosylation enzymes involved in cell membrane/envelope biogenesis (Supplementary Figure S7). However, our chemical data shows that phosphonates are associated with a high molecular weight protein fraction. We propose that in *Prochlorococcus* SB phosphonates serve as molecular decorations of glycan chains bound to high molecular membrane-anchored proteins. In glycoproteins sugar polymer chains are covalently attached to amino acid side-chains of a parent protein through post-translational modification.

Glycoproteins were once thought to be exclusive to eukaryotes, but recent efforts indicate that they are ubiquitous in bacteria and archaea[16]. In bacteria glycans are attached to specific target proteins via both N- and O-linked glycosylation[17]. Protein substrates for glycosylation reactions include the motor protein flagellins, pilins found in bacterial pilus structures[17], and S-layers which are two-dimensional paracrystalline structures that cover the outer surface of bacteria and archaea[18]. Protein glycosylation occurs via two distinct mechanisms: the direct attachment of glycan monomers or small chains to proteins via glycosyltransferases (common in flagellins and adhesins) and the pre assembly of glycan onto lipid carriers followed by transfer en masse to a protein acceptor[19]. Recently, it has been discovered that many bacteria also possess general O-linked glycosylation systems capable of promiscuous modification of numerous different types of membrane associated proteins[20–22].

In *Prochlorococcus* SB, we find genes with homologies to both S-layer biosynthesis and the general O-linked glycosylation system. S-layer biosynthetic clusters encode genes for both biosynthesis and export of lipopolysaccharide anchors[18] and the biosynthesis and export of an extracellular glycoprotein containing the critical $Ca^{2+}$- binding hemolysin domain[23,24]. In the *Prochlorococcus* SB genome between 1.15 - 1.17 Mbp we find genes encoding putative surface antigens of the filamentous hemagglutinin-glycoprotein family, $Ca^{2+}$- binding hemolysin domain proteins, and a type-I secretion system ABC transporter, and other antigen-polysaccharide repeats (Supplementary Table S2). Most compelling, however, is the presence of general O-linked glycosylation reaction enzymes adjacent to the *Prochlorococcus* SB, *Pelagibacter* HTCC7217 and *Pelagibacter* RS40 phosphonate biosynthesis clusters (Supplementary Figure S7). In *Prochlorococcus* SB we find homologs to known genes involved in extracellular capsule synthesis/export, glycan chain extension reactions, and glycosyl transferase reactions. The presence of these capsular biosynthesis enzymes is consistent with the significant cross-talk between lipopolysaccharide biosynthesis and glycoprotein biosynthesis[19] and the shared glycan substrates and enzymes between both pathways[25]. Most importantly we identify an initiating glycotransferase (PglC) which links the first glycan subunit to a lipid carrier, a flippase (Wzx) which translocates the assembled glycan-bound lipid carrier to the periplasmic face, and an O-oligosaccharyltransferase (PglL) which is the critical enzyme moving the assembled glycan chain to the final acceptor protein[25,26]. The striking mutual exclusivity we observe between phosphonate production and consumption genes in marine genomes may also be related to the molecular mechanisms and subcellular location of the machinery from both pathways. For example, both the catalytic domains of C-P lyase[27] and the assembly of glycans[16] occur on the cytoplasmic side of the bacterial inner membrane. A cell simultaneously expressing both these systems might cannibalize its own phosphonates. Taken together our findings strongly suggest that *Prochlorococcus* SB has the genomic potential to produce surface-expressed phosphonoglycoproteins.

We also note that the low frequency of phosphonate biosynthesis components in *Prochlorococcus*, and SAR11 populations is consistent with the evolutionary mechanism of negative frequency dependent selection[28]. Negative frequency dependent selection favors the presence of rare traits in a population, and negative frequency dependent patterns are well documented for microbial surface structures including lipopolysaccharides, S-layers, O-antigens, and capsular polysaccharides[29]. Rare variation in these surface structure traits, for example slight structural modification, can facilitate bacterial evasion from phage and host immune responses[30,31]. However, once a variant becomes too abundant within a population then predators, phage or the immune system may adapt to the variant, neutralizing any conferred advantage. Thus, it is theorized that natural selection in these genes results in rapid gain and loss, which ultimately makes these genes and their variants rare in the wider population. The apparent negative-frequency dependence of phosphonate producers within

*Prochlorococcus* populations suggests a role for the gene in mediating predator or phage interactions.

**Supplementary Note S5: Robust quantification of PepM and marker genes using short metagenomic reads**

Metagenomic classification of diverse functional genes can present methodological challenges because the amount of discriminative taxonomic signal in these genes may be marginal or confined to specific regions of the gene. For example, PepM appears to be horizontally transferred both within closely related microbial groups like *Prochlorococcus* and between diverse microbial lineages). In some cases, it may be difficult to taxonomically assign a short metagenome read derived from a PepM gene because the read may map equally well to a PepM sequence from multiple taxonomic lineages. We quantified the skill with which we could classify PepM sequences derived from SAR11, *Prochlorococcus*, and all bacteria and archaea genomes using mock metagenomes of known composition.

We simulated mock metagenomes from 200 randomly sampled single cell template genomes from the approximately 13000 genomes from GORG-tropics. The 200 template genomes were sampled from all available taxonomic orders but were restricted to those genomes from each order that fell into the top 10% highest estimated completeness from checkM. Genomes estimated to be below 30% completeness were excluded. These criteria resulted in 74 SAR11 and 48 *Prochlorococcus* template genomes. We simulated 150 bp paired-end Illumina sequencing reads mock metagenomes using randomreads.sh from BBMap v37.90 (https://sourceforge.net/projects/bbmap/) with 3x coverage, an insert distribution of 155-320 bp and parameters snprate=0.005, insrate=0.0005, delrate=0.0005, and subrate=0.0005. We next included all identified PepM sequences from GORG-tropics to represent the full diversity of PepM sequences in the surface ocean. As a decoy case we also included all GORG genes from the isocitrate-lyase superfamily, which have superficial similarity to PepM but lack the essential conserved active site motif "EDK(X)5NS." We then cut these authentic and decoy PepM gene sequences with 75 bp upstream and downstream flanking DNA sequence from their parent genomes and simulated sequencing reads using the same parameters for the full genomes but with a sequencing depth of 30x. This step produced sequencing reads derived from 25, 96, and 14 authentic PepM sequences from *Prochlorococcus*, SAR11, and all remaining bacteria and archaea, respectively. It also generated reads from 49 and 134 isocitrate lyase decoy sequences from bacteria and archaea and SAR11, respectively. The final resulting simulated metagenomic reads were preprocessed and merged as the Tara ocean and bioGEOTRACES metagenomes.

Because we knew the precise quantity of PepM-derived reads in the simulated metagenome we could then iteratively calculate the true positive rate, true negative rate, positive predictive value, and the F1 score (harmonic mean of the true positive rate and positive predictive value) for different combinations of sequence similarity cutoffs. In this case the positive predictive value is the probability that a metagenomic read classified as PepM is derived from a PepM sequence, the true positive rate is the percentage of true PepM reads correctly identified as such, and the true negative rate is the percentage of false PepM reads correctly identified. We selected the percent identity and E value classification cutoffs that produced the highest F1 score for PepM. Ultimately, we selected an E value cutoff of 1e-5 and a percent identity cutoff of 55%, which resulted in a true positive rate of 95% and a true negative rate of 99% for classifying any bacterioplankton PepM (Supplementary Figure S9). For *Prochlorococcus* PepM and SAR11 PepM sequences, respectively, our classification criteria produced a true positive rate of 96%/60% and a true negative rate of 97%/98%. These performance metrics suggest that in all cases we are unlikely to classify superficially related reads as a true PepM sequence. For *Prochlorococcus* and combined bacteria/archaea it appears that we correctly capture 95-96% of the total PepM sequence diversity in the mock metagenomes. The true positive rate for SAR11 was significantly lower (60%) which suggests that 40% of SAR11 PepM sequences are missed and probably assigned only to the bacteria/archaea domain level. This result is consistent with the high sequence similarity between a handful of PepM genes from SAR11 and other diverse bacterial groups, and suggests that our metagenomic database probably underestimates the contribution of SAR11 to the total inventory of PepM genes in the ocean. It probably also accounts for some of the significant positive correlation between SAR11 PepM abundance and combined bacteria/archaea PepM abundance (Supplementary Figure S5). Overall, these classification metrics indicate that we are unlikely to misclassify PepM reads from marine metagenomes and suggest our relative abundance estimates are robust.

**Supplementary Note S6: Random forest regression**

We used random forest regression, a nonparametric machine learning approach, to identify biotic and abiotic factors structuring the distributions of phosphonate producers. We selected random forest regression because the dataset has nearly 45 abiotic/biotic variables, many of which are highly correlated, and random forest is capable of handling many predictor variables and is robust to covariate co-linearities. During the cross-validation and tuning phase of the random forest model generation, we performed feature selection heuristics (see methods) to find the top 50th percentile of informative variables for *Prochlorococcus*, SAR11, and combined bacteria/archaea. We used a forest size of 1000 trees for all models. For *Prochlorococcus* we ultimately selected 10 model covariates, a minimal node size of 2, and 7 randomly sampled variables at each split resulting in a coefficient of determination ($R^2_{Pro}$) of 0.74 and Root Mean Square Error ($RMSE_{Pro}$) of 0.04. For SAR11 we selected 15 selected model covariates, a minimal node size of 2, and 6 randomly sampled variables at each split resulting in a coefficient of determination ($R^2_{SAR11}$) of 0.69 and Root Mean Square Error ($RMSE_{SAR11}$) of 0.03. For the combined bacteria/archaea we selected 21 selected model covariates, a minimal node size of 2, and 8 randomly sampled variables at each split resulting in a coefficient of determination ($R^2_{combined}$) of 0.94 and Root Mean Square Error ($RMSE_{combined}$) of 0.02. We then assessed whether these top 50th percentile variables were more informative to the model than including random information, using the Boruta heuristic[32]. The Boruta algorithm is a robust, statistically grounded feature ranking method that aims to identify all variables in a dataset that contain useful information for model predictive outcome. This approach is useful when the variables driving the performance model are of interest and not necessarily the ultimate predictive outcome itself. Boruta performs feature ranking by comparing all variables with randomized versions of themselves in an iterative framework. In addition to variable ranking based on relative scores, Boruta, therefore, provides a measure of the significance of each variable relative to noise.

For *Prochlorococcus* and SAR11 there was an appreciable amount of variation in producers that the models could not capture ($R^2_{Pro}$= 0.74, $R^2_{SAR11}$= 0.69). There were 10 and 15 covariates that showed better predictive utility than randomly shuffled data for *Prochlorococcus* and SAR11, respectively (Figure S3). However, most predictive utility was concentrated into the top 5 variables, while the importance of the other lower ranking variables was in most cases only marginally higher than noise. The five most highly predictive variables for *Prochlorococcus* phosphonate producers (Supplementary Figure S3 and S5, Supplementary Table S1) included inorganic phosphate concentrations, the modeled climatological mean dissolved organic phosphorus concentrations, the subsurface chlorophyll maximum layer type, and the relative abundance of the *Prochorococcus* LLIV clade. The subsurface chlorophyll maximum layer typestypes 1-4 were characterized by deepening SCML depths (~ 20 to 115 meters) while types 5 and 6 were characterized by high fluorescence throughout the upper 75 meters. The top five variables for SAR11 were the relative abundance of total phosphonate producers in the sample, the climatological mean nitrate concentrations, SAR11 IV relative abundance, SAR11 clade IIb relative abundance, and dissolved aluminum concentrations (Supplementary Figure S3 and S5, Supplementary Table S1). The integrated bacteria/archaea phosphonate producer model performed best ($R^2_{combined}$ = 0.94) and the top five variables (SAR11 clade IIb relative abundance, SAR11 PepM relative abundance, archaea relative abundance, SAR11 clade IV relative abundance, Depth) were strongly informative for the random forest prediction when compared to noise. We also confirmed these factors for all groups using parametric beta-binomial regression (Supplementary Table S1)[33].

### Supplementary Tables

#### Table S1: Model coefficients from beta-binomial regression of PepM abundance

| Taxonomic group | Term | Estimate | 95% CI [lower] | 95% CI [upper] | Std. Error | t value | Bootstrapped LRT P value | LRT P value | Term Significance |
|---|---|---|---|---|---|---|---|---|---|
| Prochlorococcus | *Prochlorococcus* [%] | -0.011 | -0.015 | -0.007 | 0.003 | -4.433 | 0.004 | 4.05E-34 | *** |
| Prochlorococcus | Pro LLIV [%] | 0.011 | 0.002 | 0.018 | 0.006 | 1.771 | 0.077 | 6.11E-02 | . |
| Prochlorococcus | DOP Clim. avg [umol kg-1] | 1.342 | 0.360 | 2.385 | 0.567 | 2.366 | 0.009 | 4.53E-04 | * |
| Prochlorococcus | PO4 [umol kg-1] | -0.014 | -0.197 | 0.167 | 0.126 | -0.107 | 0.745 | 7.13E-01 | NS |
| Prochlorococcus | DCM type 2 | -0.164 | -0.304 | -0.043 | 0.077 | -2.129 | 0.003 | 6.41E-22 | * |
| Prochlorococcus | DCM type 3 | -0.561 | -0.710 | -0.419 | 0.085 | -6.567 | 0.003 | 6.41E-22 | *** |
| Prochlorococcus | DCM type 4 | -0.250 | -0.380 | -0.125 | 0.076 | -3.277 | 0.003 | 6.41E-22 | ** |
| Prochlorococcus | DCM type 5 | -0.721 | -0.970 | -0.473 | 0.134 | -5.387 | 0.003 | 6.41E-22 | *** |
| Prochlorococcus | DCM type 6 | -0.084 | -0.210 | 0.037 | 0.074 | -1.136 | 0.003 | 6.41E-22 | NS |
| SAR11 | Total PepM [per genome] | 0.021 | 0.018 | 0.024 | 0.002 | 11.502 | 0.001 | 6.94E-29 | *** |
| SAR11 | NO3 Clim. avg [umol kg-1] | 0.004 | 0.001 | 0.006 | 0.001 | 2.640 | 0.120 | 2.95E-04 | ** |
| SAR11 | SAR11 IV [%] | -0.014 | -0.021 | -0.007 | 0.004 | -3.908 | 0.062 | 3.34E-05 | *** |
| SAR11 | SAR11 IIb [%] | -0.005 | -0.007 | -0.001 | 0.002 | -2.498 | 0.003 | 5.99E-05 | * |
| SAR11 | Dissolved Aluminum [nmol kg-1] | 0.003 | 0.001 | 0.004 | 0.001 | 3.370 | 0.003 | 1.92E-04 | *** |
| All bacterioplankton | SAR11 IIb [%] | 0.0114 | 0.0088 | 0.0137 | 0.0017 | 6.835 | < 0.001 | 6.08E-13 | *** |
| All bacterioplankton | SAR11 PepM [per genome] | 0.0452 | 0.0394 | 0.0506 | 0.0035 | 12.901 | < 0.001 | 8.93E-34 | *** |
| All bacterioplankton | Archaea [%] | 0.0198 | 0.0165 | 0.0223 | 0.0022 | 9.083 | < 0.001 | 1.70E-21 | *** |
| All bacterioplankton | SAR11 IV [%] | -0.0148 | -0.0209 | -0.0099 | 0.0035 | -4.237 | < 0.001 | 2.31E-08 | *** |
| All bacterioplankton | Pro LLIV [%] | 0.0047 | 0.0031 | 0.0066 | 0.0016 | 2.902 | < 0.001 | 1.88E-07 | ** |
| All bacterioplankton | Depth [m] | 0.0006 | 0.0004 | 0.0008 | 0.0001 | 4.117 | < 0.001 | 1.88E-07 | *** |

Coefficient estimates are of unit log-odd change in the probability of sampling a PepM read due to a one unit covariate change while controlling for the other covariates. For the categorical variable DCM type the coefficient estimate is from the reference level DCM type 1. 95% confidence intervals are simulated from 1000 random draws from the beta-binomial distribution parameterized from the covariate data. Each covariate is tested for statistical significance using a parametric bootstrapped likelihood ratio test (N=1000) and a classic likelihood ratio (LRT) test using a canonical Chi-squared distribution. Term significance: *** P < 0.001, ** 0.001 < P < 0.01, * 0.01 < P < 0.05, . 0.05 < P < 0.1, NS P > 0.1. *Prochlorococcus* and Archaea are percentage of all successfully mapped reads to MARMICRODB. Clades LLIV, SAR11 IV, SAR11 IIb are percentage of reads mapped to *Prochlorococcus* and SAR11, respectively. Clim. avg, climatological average from the MIT Darwin model; DCM depth range, deep chlorophyll maximum layer depth category – 1=shallow, 6=deep.

**Table S2: Functional enrichment of polysaccharide biosynthesis genes within phosphonate biosynthesis gene clusters.**

| Genome | Kegg Pathway | Gene Ratio | Background Ratio | p.adjust | qvalue | KO | Gene function Description |
|---|---|---|---|---|---|---|---|
| SB | map00520 | 6/16 | 24/754 | 7×10-5 | 6×10-5 | K02377, K02377 | Nad-dependent epimerase dehydratase |
| SB | map00520 | 6/16 | 24/754 | 7×10-5 | 6×10-5 | K01784 | udp-glucose 4-epimerase |
| SB | map00520 | 6/16 | 24/754 | 7×10-5 | 6×10-5 | K01710, K08679 | Nucleotide sugar epimerase |
| SB | map00520 | 6/16 | 24/754 | 7×10-5 | 6×10-5 | K01711 | Gdp-mannose 4,6-dehydratase |
| SB | map00520 | 6/16 | 24/754 | 7×10-5 | 6×10-5 | K00971, K16011 | Mannose-1-phosphate guanylyltransferase |
| SB | map00520 | 6/16 | 24/754 | 7×10-5 | 6×10-5 | K00012 | Udp-glucose 6-dehydrogenase |
| RS40 | map00520 | 6/28 | 29/755 | 1×10-2 | 1×10-2 | K01784 | epimerase dehydratase |
| RS40 | map00520 | 6/28 | 29/755 | 1×10-2 | 1×10-2 | K13010 | DegT/DnrJ/EryC1/StrS aminotransferase family |
| RS40 | map00520 | 6/28 | 29/755 | 1×10-2 | 1×10-2 | K01654, K18430 | synthase |
| RS40 | map00520 | 6/28 | 29/755 | 1×10-2 | 1×10-2 | K01791, K18429 | UDP-N-acetylglucosamine 2-epimerase |
| RS40 | map00520 | 6/28 | 29/755 | 1×10-2 | 1×10-2 | K00966, K16881 | Glucose-1-phosphate cytidylyltransferase |
| RS40 | map00520 | 6/28 | 29/755 | 1×10-2 | 1×10-2 | K01654, K18430 | synthase |

Subset of results from a Universal Enrichment Analysis implemented with the bioconductor package clusterprofiler. The remainder of the results for *Prochlorococcus* and SAR11 are available from https://github.com/slhogle/phosphonates. SB, *Prochlorococcus* sp. SB; RS40, *Candidatus Pelagibacter* sp. RS40; map00520, Amino sugar and nucleotide sugar metabolism; KO, Kegg Orthology identifier;

p.adjust, Bounded False Discovery Rate (FDR) adjusted probability for hypergeometric test statistic under the null hypothesis; qvalue, direct estimate of the FDR associated with p.adjust; Gene Ratio, k/n where n is the total number of genes within 10000 nucleotides upstream/downstream of the PepM gene and k is the number of genes in that region annotated to the Kegg Pathway; Background ratio, M/N where N is the total number of genes with a Kegg Orthology assignment and M is the number of Kegg Orthology assigned genes annotated to the Kegg Pathway.

**Table S3: *Prochlorococcus* SB phosphonate (Phn) and phosphate (Ph) cellular proportions.**

| Harvest stage | # | Phn proportion | Ph proportion |
|---|---|---|---|
| *Prochlorocccus* SB P-replete (exponential phase) | 1 | 0.41 | 0.59 |
| | 2 | 0.43 | 0.57 |
| | mean | 0.42 | 0.58 |
| *Prochlorocccus* SB P-limited (stationary phase) | 1 | 0.70 | 0.30 |
| | 2 | 0.71 | 0.29 |
| | mean | 0.71 | 0.29 |

$^{31}$P-NMR spectra of each conditions and each replicates were manually integrated and those values used to calculate the proportion of phosphonate and phosphate in the cells.

**Table S4: *Prochlorococcus* SB phosphorus (P), phosphonate (Phn) and phosphate (Ph) to carbon ratios.**

| Harvest stage | # | P/C | Phn/C | Ph/C |
|---|---|---|---|---|
| *Prochlorocccus* SB P-replete (exponential phase) | 1 | 0.009 ± 0.001 | 0.0036 ± 0.0005 | 0.0052 ± 0.0007 |
| | 2 | 0.0057 ± 0.0005 | 0.0024 ± 0.0002 | 0.0033 ± 0.0003 |
| | mean | 0.007 ± 0.001 | 0.0030 ± 0.0004 | 0.0042 ± 0.0007 |
| *Prochlorocccus* SB P-limited (stationary phase) | 1 | 0.008 ± 0.001 | 0.0059 ± 0.0009 | 0.0025 ± 0.0004 |
| | 2 | 0.0070 ± 0.0002 | 0.0050 ± 0.0002 | 0.00203 ± 0.00007 |
| | mean | 0.0076 ± 0.0005 | 0.0054 ± 0.0003 | 0.0047 ± 0.0002 |

The proportions of phosphonate and phosphate in the cells and the P/C ratio are used to calculate the Phn/C and C/Ph ratios. Errors associated the P/C ratio are calculated using the errors associated with the %C and %P in the cells and as: $\Delta\frac{P}{C} = \frac{P}{C} \times \sqrt{(\frac{\Delta\%C}{\%C})^2 + (\frac{\Delta\%P}{\%P})^2}$. Errors associated with Phn/C and Ph/C ratios are calculated as: $\Delta\frac{Phn}{C} = \frac{Phn}{C} \times \sqrt{(\frac{\Delta P/C}{P/C})^2}$. Errors associated with the mean values are calculated using the standard error of the mean.

**Table S5: Comparison of genome, PepM, and random tree topologies.**

| Tree | RF | Max RF | Norm RF | Eff tree size | Ref in Src | Src in Ref |
|---|---|---|---|---|---|---|
| Gphylo vs Gphylo | 0 | 1984 | 0 | 999 | 1 | 1 |
| PepM vs Gphylo | 1372 | 1926 | 0.71 | 999 | 0.64 | 0.66 |
| Rtrees vs Gphylo (n=15) | 1708 ± 18 | 1988 | 0.86 ± 0.01 | 999 | 0.570 ± 0.005 | 0.570 ± 0.005 |
| Gphylo vs Gphylo | 0 | 26 | 0 | 16 | 1 | 1 |
| PepM vs Gphylo | 22 | 26 | 0.85 | 16 | 0.61 | 0.61 |
| Rtrees vs Gphylo (n=15) | 21.8 ± 1.9 | 26 | 0.84 ± 0.08 | 16 | 0.61 ± 0.04 | 0.61 ± 0.04 |

When compared to the concatenated genome phylogeny (Gphylo), 10 trees simulated at random (Rtrees) have similar Robinson-Foulds distances to the PepM phylogeny (PepM) indicating that the PepM evolutionary history corresponds poorly to that of the core genome in both archaea and bacteria. The upper half of the table compared the bacterial genome phylogeny while the lower half compares the archaeal genome phylogeny. Column descriptions: Tree, tree comparison performed; RF, calculated Robinson-Foulds metric; Max RF, maximum possible RF distance for the comparison; Norm RF, for normalized comparison between different tree comparisons (RF/Max RF); Eff tree size, effective tree size used for the calculation after pruning unshared leaves; Ref in Src, normalized number of splits from Gphylo found in either PepM or Rtrees; Src in Ref, normalized number of splits from either PepM or Rtrees found in Gphylo.

### Supplementary Figures

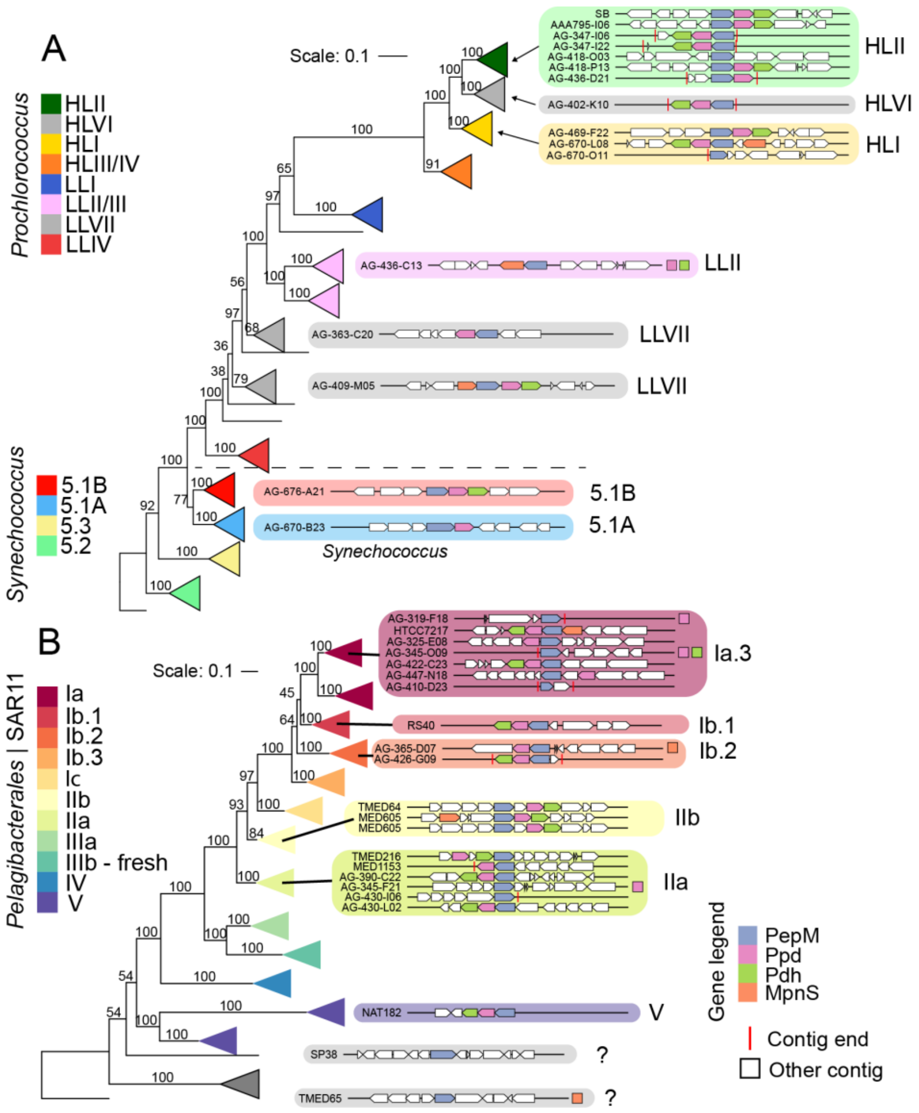


**Figure S1: A) Prochlorococcus and B) SAR11 phosphonate producers**

Phylogenies are constructed from 120 concatenated, single-copy marker genes from *Prochlorococcus* (112 genomes) and *Synechococcus* (23 genomes) and SAR11 (92 genomes). Scale bar is 0.1 amino acid substitutions. The cyanobacteria tree is rooted at *Synechococcus* sp. WH5701 and the SAR11 tree is rooted at TMED13, a metagenome-assembled genome from Tara Oceans distantly related to 'Candidatus Pelagibacter ubique.' Bootstrap values are based on 250 resamplings. Monophyletic ecotypes are represented by colored cartoon triangles. Inclusion of new genomes in this study resulted in a polyphyletic origin for the original SAR11 clade Ib, therefore we have partitioned clade Ib into 3 monophyletic subclades (Ib.1, Ib.2 and Ib.3). Phosphonate biosynthetic gene cluster diagrams for all genomes from each ecotype are displayed to the right. Vertical red lines represent contig breaks in the gene diagrams and filled squares next to the diagram indicate that the gene was detected on a separate noncontiguous contig. For simplification, MpnS category also includes the functionally related enzyme HepDI.

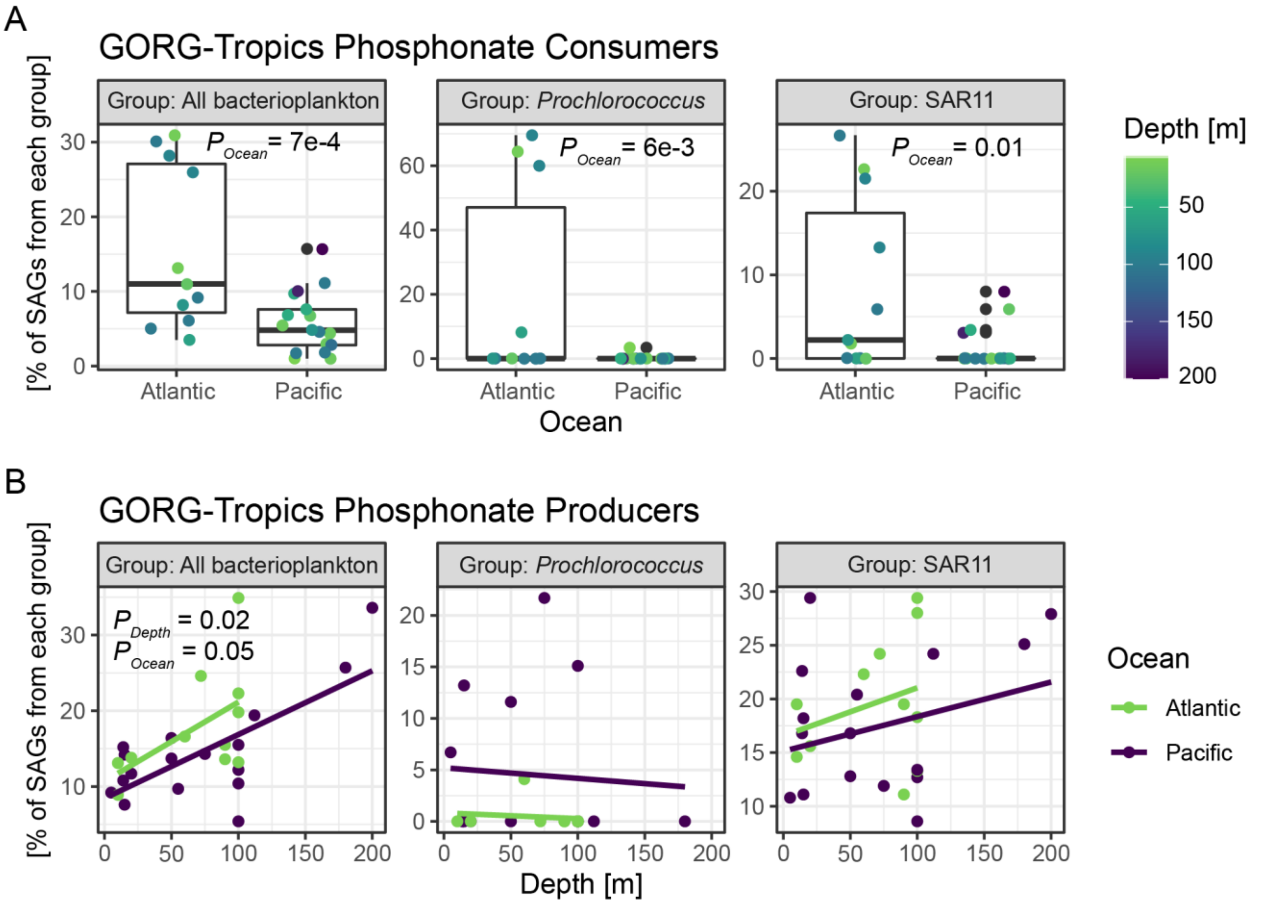


**Figure S2 Phosphonate consumers and producers in 28 GORG-Tropics samples**

Same as in Fig 2C,D but includes *Prochlorococcus* and SAR11. Proportions [%] are total producers or consumers divided by the total number GORG assemblies and are corrected using the estimated sequence recovery from assemblies (see methods). P values for covariates in beta-binomial regression are displayed if statistically significant ($P <= 0.05$). Beta-binomial regression all consumers; Ocean - Est=-1.04, Err=0.27, t=-3.81, P=7e-4; link=logit; log L = -80.425, df=4, resid df=24. Beta-binomial regression *Prochlorococcus* consumers; Ocean - Est=-3.50, Err=1.15, t=-3.04, P=6e-3; link=logit; log L = -20.181, df=4, resid df=21. Beta-binomial regression SAR11 consumers; Ocean - Est=-1.73, Err=0.65, t=-2.66, P=0.01; link=logit; log L = -41.916, df=4, resid df=24. Beta-binomial regression all producers; Depth - Est=0.0036, Err=0.0014, t=2.604, P=0.02; Ocean - Est=-0.30, Err=0.14, t=-2.08, P=0.05; link=logit; log L=-78.179, df=4, resid df=24.

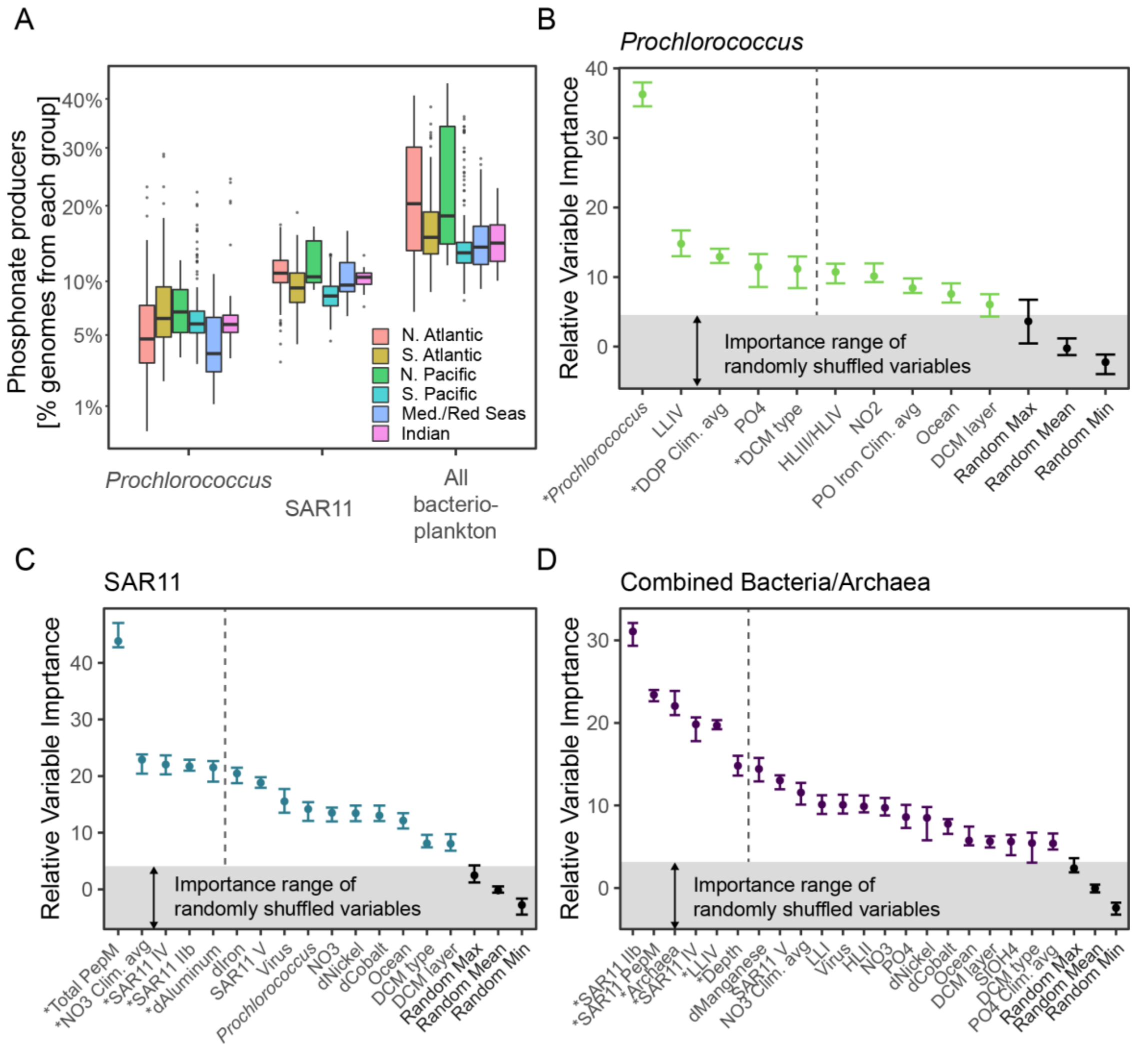


**Figure S3: Environmental factors driving the relative abundance of PepM in *Prochlorococcus*, SAR11, and combined Bacteria/Archaea**

Same as in Figure 3A but binned by ocean region. There is no statistically significant difference between ocean regions for *Prochlorococcus*, SAR11, or combined Bacteria/Archaea PepM relative abundance. Black line is the median value, the box shows the 25th and 75th percentiles, and the whiskers extend to 1.5 of the interquartile range for n metagenome samples in each group. Individual points are outliers, and the vertical axis is square-root transformed. B-D) Random forest variable importance measures for each taxonomic group. Point ranges show median and maximum/minimum for 100 random permutations of the variable importance algorithm. Only informative variables for predicting PepM distributions are displayed. Randomized controls in black represent the distribution of the median, minimum, and maximum importance of randomly shuffled variables at each algorithm iteration and can be interpreted as the baseline variable importance expected simply due to chance. Asterisks indicate whether a variable was also statistically significant in a parametric beta-binomial regression (Supplementary Table S1). The top five predictive variables are demarcated by a dashed line. *Prochlorococcus*, Archaea, and viruses are units percentage of all successfully mapped reads to MARMICRODB. Clades HLII, LLI, LLIV, HLIII/HLIV, SAR11 IV, SAR11 IIb, SAR11 V, are percentage of reads mapped to *Prochlorococcus* and SAR11, respectively. Clim. avg, climatological average from the MIT Darwin model; DCM depth range, deep chlorophyll maximum layer depth category – 1=shallow, 6=deep. PepM units are the percentage of *Prochlorococcus*, SAR11, or all genomes with the trait.

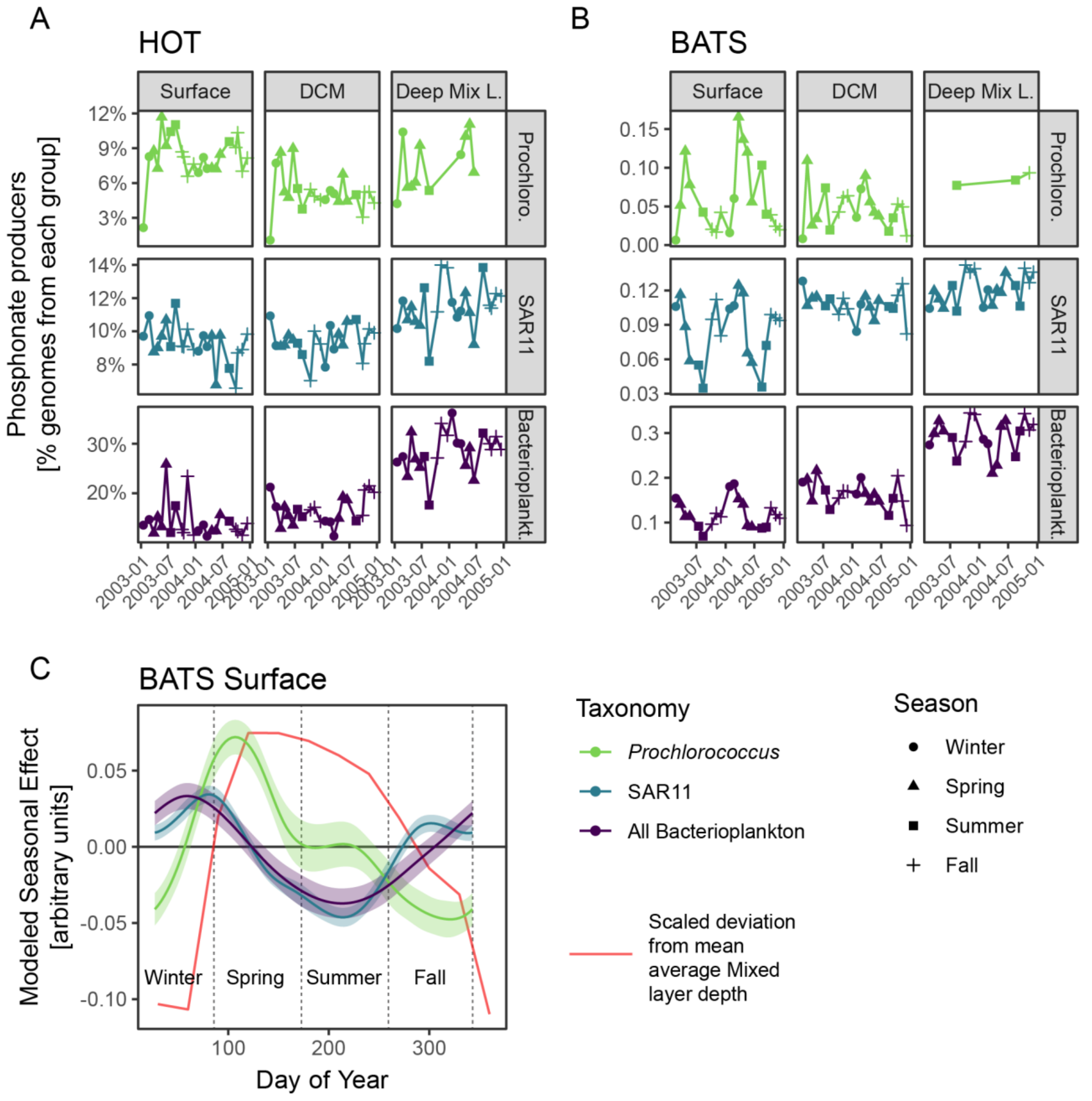


**Figure S4: Time series of PepM relative abundance at the Hawaii Ocean Time-series (HOT) and the Bermuda Atlantic Time Series (BATS).**

PepM relative abundance shown by depth range (horizontal grid axis, DCM = deep chlorophyll maximum layer, Deep Mix L. = deep mixed layer, 1% photosynthetically active radiation) and by taxonomic grouping (vertical grid) at **A)** HOT and **B)** BATS. Horizontal axis shows time and points are shaped by sampling season. **C)** The modeled seasonal effect at BATS surface on the relative abundance of PepM of each taxonomic grouping. [GAM SAR11, family=Gamma, link=identity: s(dayOfYear), edf=5.4, Ref. df=8, F=9.8, P=7e-5. GAM Prochlorococcus, family=Gamma, link=identity: s(dayOfYearDCM), edf=3.8, Ref. df=8, F=4.5, P=0.001, GAM Bacterioplankton, family=Gamma, link=identity: s(dayOfYearDCM), edf=3.2, Ref. df=8, F=5.0, P=2e-4]. The red line shows the scaled deviation from the annual mean of the mixed layer depth at BATS for the time range in **B)**. For example, a negative deviation indicates that the mixed layer depth is deeper than the annual mean.

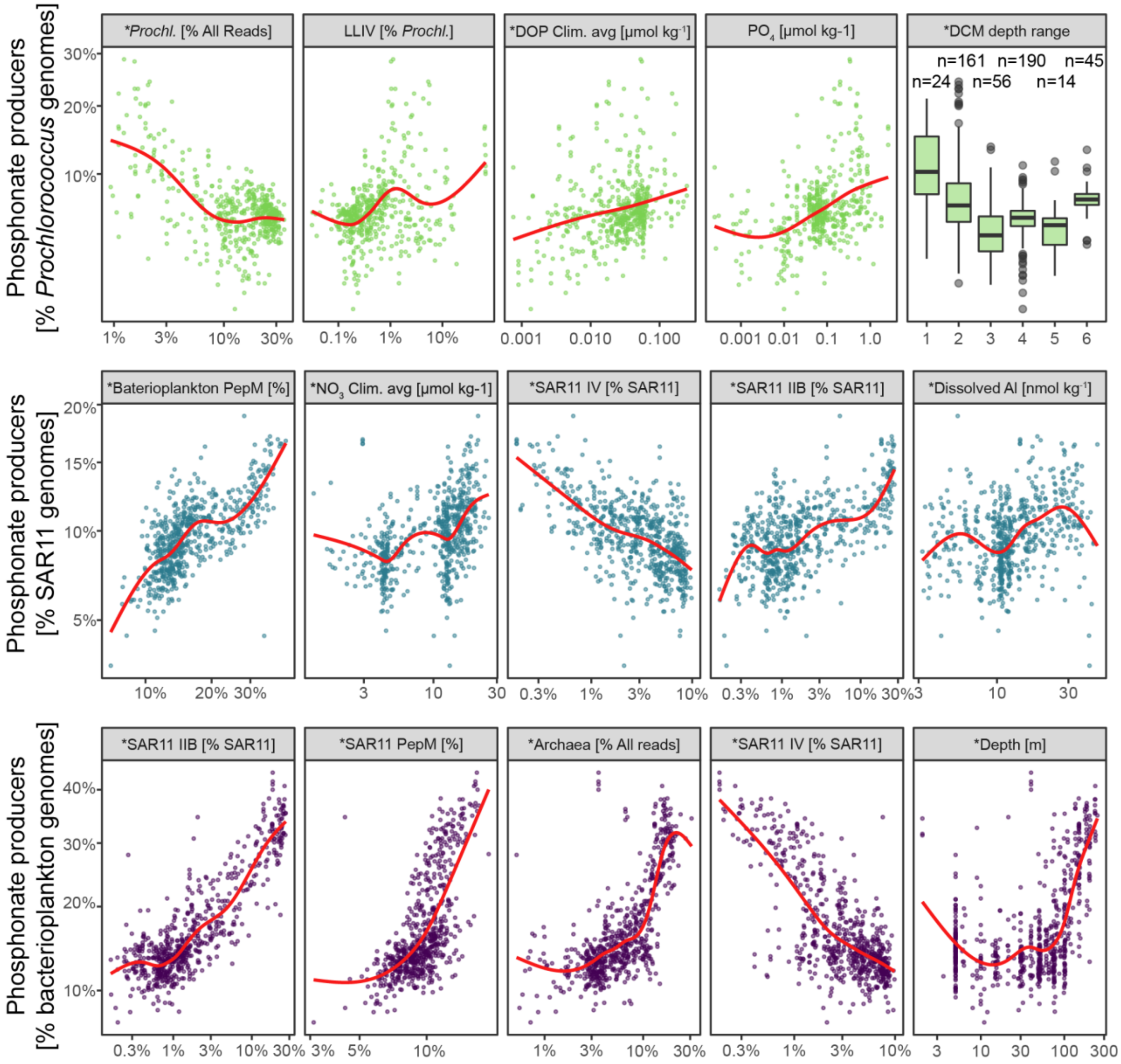


**Figure S5: Relationship between *Prochlorococcus*, SAR11, and combined Bacteria/Archaea PepM relative abundance and the top five biotic/abiotic covariates for each taxonomic group.**

Phosphonate producer relative abundance (vertical axis) vs environmental features (horizontal axis) for the top five most informative variables from random forest regressions of each taxonomic group (Figure 3). Each point is a metagenomic observation. Climatological averages are from the MIT Darwin model while the other biotic/abiotic variables are in situ measurements from GEOTRACES and Tara Oceans. Red lines are local polynomial regression fits to the data. Asterisks indicate whether a variable was also statistically significant in a parametric beta-binomial regression (Supplementary Table S1). *Prochlorococcus* and Archaea are percentage of all successfully mapped reads to MARMICRODB. Clades LLIV, SAR11 IV, SAR11 IIb are percentage of reads mapped to *Prochlorococcus* and SAR11, respectively. Clim. avg, climatological average from the MIT Darwin model; DCM depth range, deep chlorophyll maximum layer depth category – 1=shallow, 6=deep. PepM units are the percentage of *Prochlorococcus*, SAR11, or all genomes with the trait.

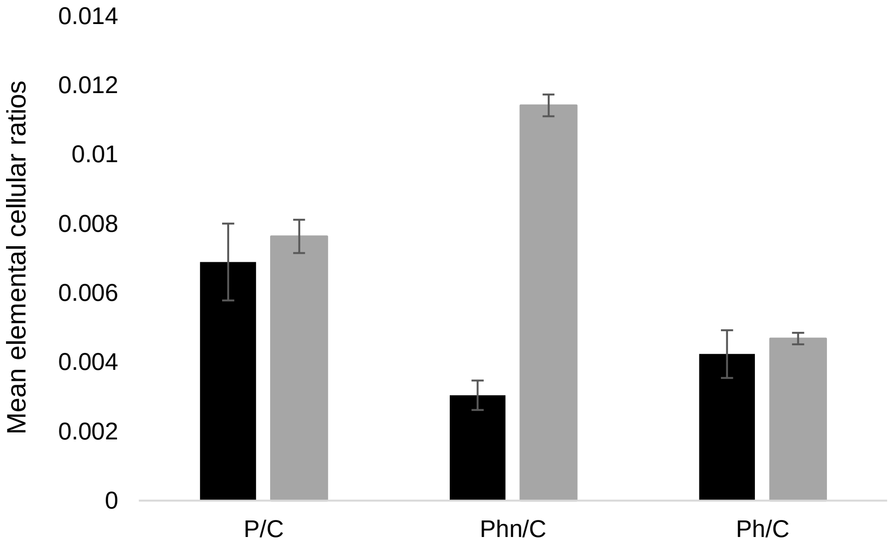


**Figure S6: Variations of *Prochlorococcus* SB P/C, Phn/C and Ph/C ratios between in exponential (P-replete; black) and stationary phase (P-limited; grey).** When *Prochlorococcus* SB cells become P-limited, their P/C ratio increases slightly whereas their Phn/C ratio increases and their Ph/C ratio decreases indicating an increase in phosphonate content and a decrease in phosphate (Supplementary Table S4). As before, errors associated with the mean values are calculated using the standard error of the mean.

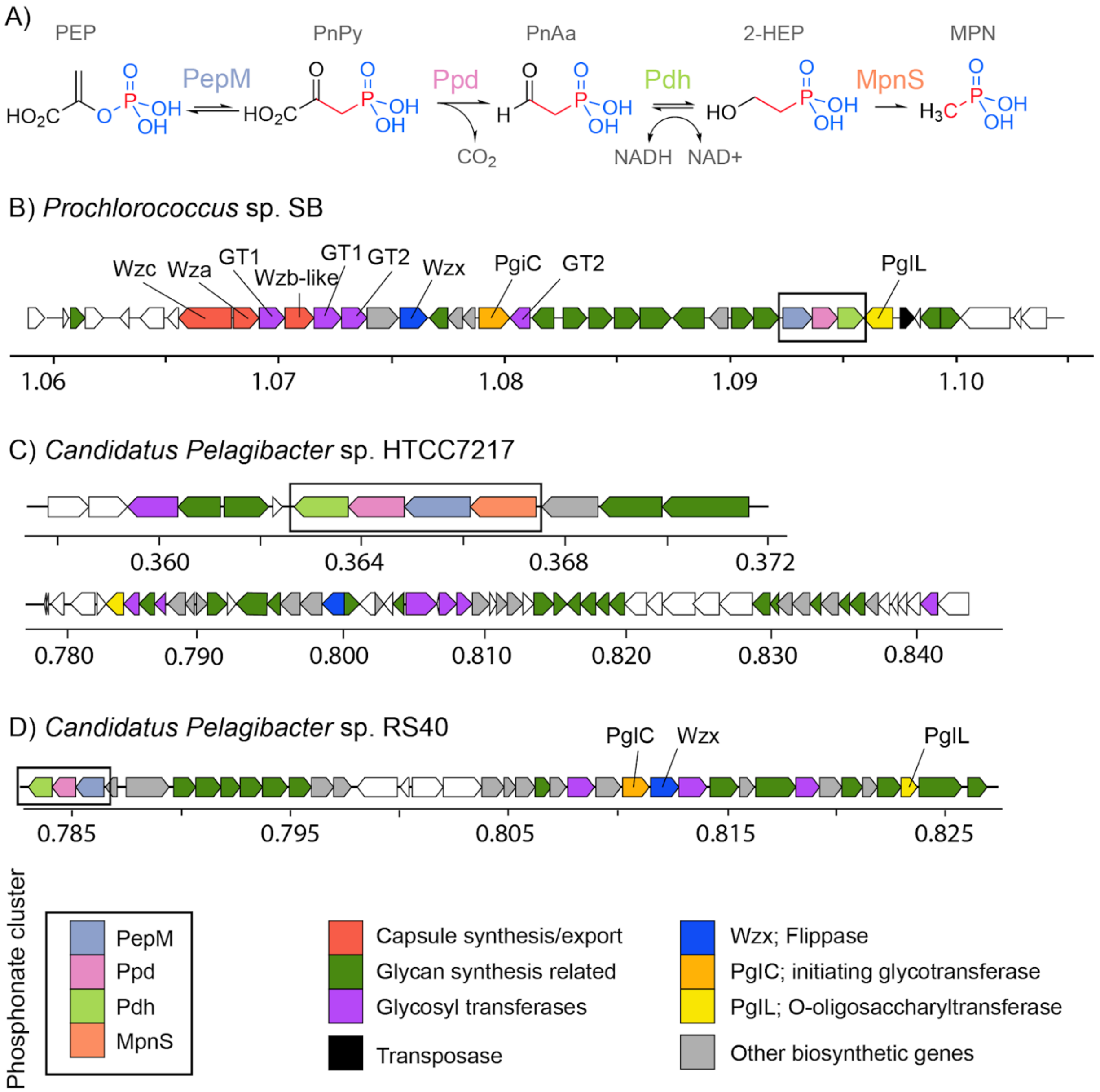


**Figure S7: Polysaccharide and phosphonate biosynthesis in *Prochlorococcus* and SAR11 isolate genomes.**

**A)** same reaction diagram as in Figure 1 with chemical intermediates (black) and catalyzing enzymes (color) **B-D)** Putative phosphonoglycoprotein biosynthesis gene clusters in *Prochlorococcus* SB, *Candidatus Pelagibacter* sp. HTCC7217, and *Candidatus Pelagibacter* sp. RS40 genomes annotated by antiSMASH[34] and eggNOG mapper[35]. Genome coordinates are $\times 10^6$ base pairs. Phosphonate biosynthetic genes are colored by enzyme as in **A)** and are shown in black boxes. Genes involved in modular biosynthesis of sugars are shown in red, green, and purple. PgiC is the putative initiating glycotransferase, which links the first glycan subunit to a lipid carrier, Wzx is a flippase which translocates the assembled glycan-bound lipid carrier to the periplasmic face, and PglL is an O-oligosaccharyltransferase which is the critical enzyme moving the assembled glycan chain to the final acceptor protein. Other biosynthetic genes (grey) have functional predictions not directly related to glycan biosynthesis.

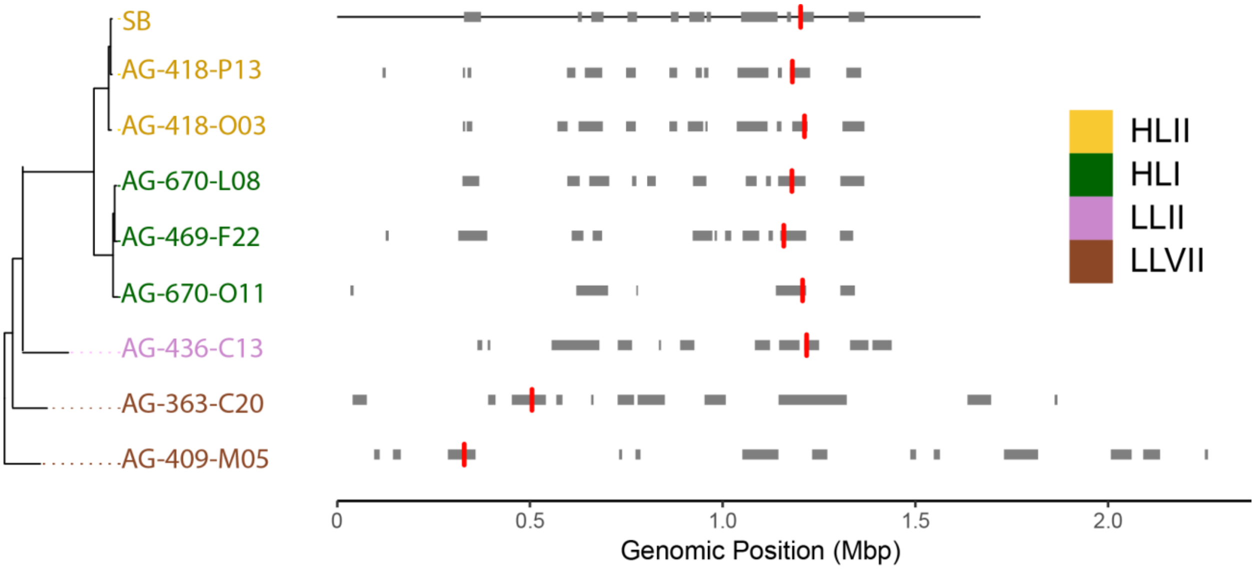


**Figure S8: Distribution of phosphonate biosynthesis gene clusters within *Prochlorococcus* genomic islands.**

Includes a subset of all *Prochlorococcus* genomes with i) high enough completeness and low enough contig fragmentation for island prediction (see methods) and ii) that contain phosphonate biosynthesis genes. Genomes are ordered by phylogeny (left) and colored by ecotype/clade (right). Grey bars denote the location of predicted genomic islands and red vertical lines denote the location of phosphonate biosynthesis clusters within predicted genomic islands. *Prochlorococcus* SB genome assembly length is shown as a black horizontal line. The assembly lengths of single cell genomes are omitted due to incompleteness and contig fragmentation. Single cell genome contigs and genomic islands are arbitrarily ordered to *Prochlorococcus* SB.

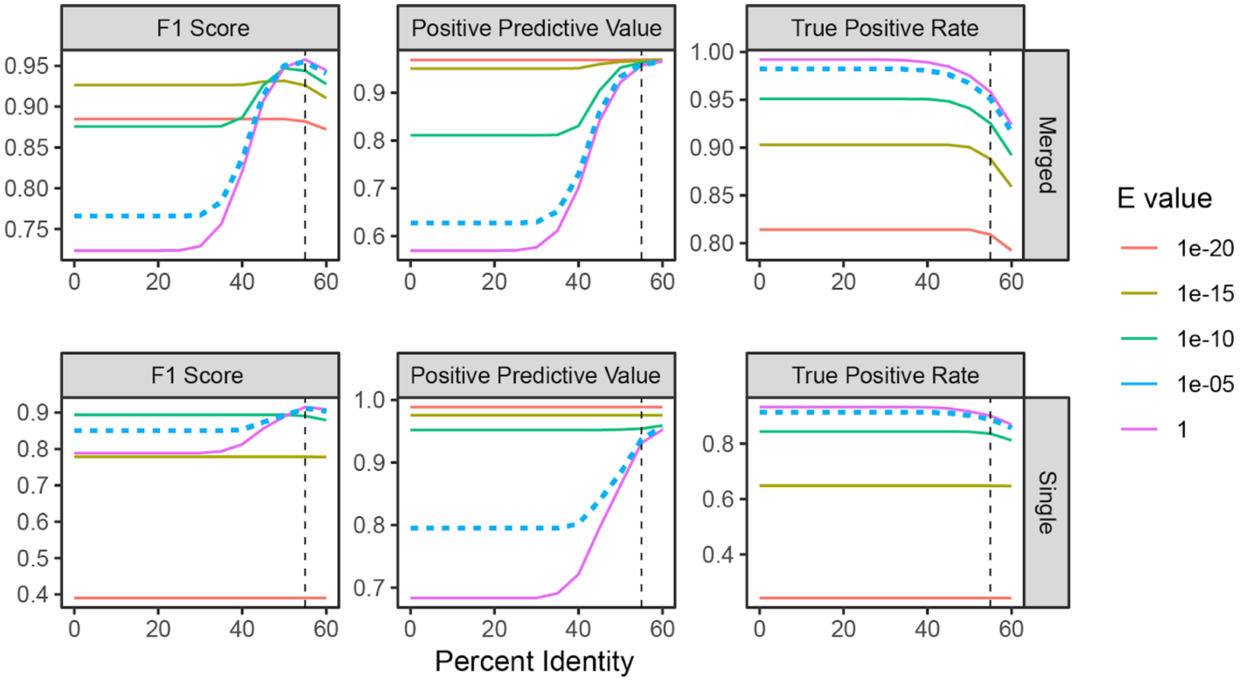


**Figure S9: Empirical determination of similarity cutoffs for identifying PepM-derived metagenomic short sequencing reads.**

F1 score, Positive Predictive Value, and True Positive Rate for classifying PepM from mock metagenomes simulated from 150 bp Illumina paired-end reads. Results are shown both for merged read pairs (top), and the best scoring read from an unmerged read pair (bottom). The True Postive Rate measures the proportion of “true” simulated PepM reads correctly identified by a E value/Percent Identity cutoff combination while Positive Predictive Value measures the proportion of “true” simulated PepM reads that are correctly identified out of all reads passing the E value/Percent Identity cutoff combination. The F1 score is the harmonic mean of precision and sensitivity. Cutoffs were empirically chosen to maximize the F1 score resulting in an E value threshold of 1e-5 (dashed blue line) and a Percent Identity cutoff of 55% (dashed vertical line).
